# Transformation and recombination of neural information in a brain network

**DOI:** 10.64898/2026.03.04.709604

**Authors:** Ying Jiang, Yuting Ke, Jingkai Wen, Johan Medrano, Wenyu Tu, Brenna Stallings, Sarah Bricault, Anna Dong, Sunho Kevin Chung, Alan Jasanoff

## Abstract

Mammalian brain function relies on integrated interactions among interconnected neural structures. The principles by which regional activity is transformed into projection-specific signals and then recombined at targets elsewhere in the brain are fundamental to brain processing, but remain poorly understood. Here we study these phenomena using a genetically encoded probe that provides neurophysiological readouts from virally labeled projections on a brain-wide scale via functional magnetic resonance imaging (fMRI). By analyzing outputs from thalamic and cortical somatosensory processing regions in rats, we find that projection-specific neural population activity undergoes shifts in tuning and temporal characteristics as it emanates from source regions. Patterns of neural information flow to targeted brain structures reconfigure under different conditions of stimulation and rest, contrasting with intrinsic fMRI functional connectivity profiles, which remain constant. Excitatory and inhibitory projections are coactivated during stimulation, but their relative response amplitudes change dynamically between stimulus conditions and across repeated stimuli, suggesting mechanistic roles for network-wide shifts in excitation/inhibition balance. Our results thus reveal how information flow throughout a neural system reshapes to promote stimulus selectivity and provide underpinnings of large-scale brain phenomena.

---

Understanding brain function at a systems level requires deciphering how interactions among distributed neural structures underlie perception and cognition^1–3^. Computational roles of individual brain regions depend critically on how they integrate inputs and disseminate output to distal areas. Although anatomical substrates of interregional communication are increasingly well characterized^4–6^, the propagation throughout projection systems of neural signals themselves is less understood. In humans and animals with opaque brains, network-level descriptions of brain-wide function typically rely on approaches such as functional connectivity (FC) analysis^7, 8^, in which interactions between brain regions are inferred from correlative relationships, or dynamic causal modeling^9^, which interprets multiregional neural dynamics under the simplifying hypothesis that each brain area affects other regions in proportion to its own activity. Empirical measurements of projection-specific signaling could test the assumptions of such models while directly revealing how neural activity is transferred and transformed between brain structures.

The signaling properties of brain-wide projection systems can be measured using a recently reported genetically encodable probe detectable by functional magnetic resonance imaging (fMRI)^10^. The probe, referred to as nitric oxide synthase for targeting imaging contrast (NOSTIC), is an enzyme-based neural activity sensor that employs a calmodulin-interacting domain to transduce intracellular calcium fluctuations into production of the potent vasodilator NO^11^. Neurons that express NOSTIC drive activity-dependent hemodynamic signals that are visualized with temporal resolution on the order of seconds, using the same blood oxygenation level dependent (BOLD) contrast mechanism that underlies conventional fMRI. NOSTIC-dependent hemodynamic readouts can be distinguished from intrinsic fMRI signals using a drug, 1400W^12^, that selectively suppresses NOSTIC but does not affect endogenous BOLD signals^10^. Circuit-specific fMRI is performed by by using a retrogradely transported herpes simplex viral vector^13^ (HSV-NOSTIC) to drive expression of NOSTIC in neural populations presynaptic to viral injection sites in the brain. Functional imaging with NOSTIC leverages the whole-brain spatial coverage of fMRI while revealing the specific contributions of labeled neural populations to network activity. This approach thus offers a means for measuring the flow of information among brain areas at mesoscopic scale.

The rodent somatosensory system provides a paradigmatic context for studying principles of projection-dependent signaling in the brain. Primary somatosensory cortex (S1) receives ascending input from ventroposterior thalamus (VP) and from posteromedial thalamus (POm)^14, 15^, a widely connected region which itself receives feedback from S1 and a broad set of other sources^16^. Modulatory input comes from structures including the zona incerta (ZI) and superior colliculus (SC)^17, 18^, while higher order stimulus processing takes place in secondary somatosensory cortex (S2) and further cortical areas that ultimately contribute to motor behavior^19, 20^. These brain regions have close homologs in primates^21, 22^ and constitute a networked architecture that is analogous to other sensory processing systems^23–25^. Interactions among the regions contribute to canonical senory processing features like tuning and adaptation^26^, as well as cognitive phenomena like attention and perceptual decision-making^27^, but network-level operations involved in these phenomena are still unclear. By applying NOSTIC in neural populations that project among somatosensory structures, system-wide contributions to fundamental processing mechanisms can be probed while also addressing general questions about input/output relationships in thalamocortical circuitry.

## NOSTIC reports somatosensory activity

To obtain targeted fMRI measurements from a large set of somatosensory projections, we injected an updated HSV-NOSTIC vector that encodes NOSTIC fused to the fluorescent protein mCherry (**>Supplementary Table 1** and **Supplementary Figure 1**) unilaterally into the POm nucleus of eight adult rats. Following at least three weeks of expression time, animals were anesthetized and fMRI was performed on a 7 T scanner during trains of electrical stimuli delivered to the forelimbs^28^, first before and then after intravenous injection of the NOSTIC-inhibiting drug 1400W (25 mg/kg), which spares endogenous fMRI signals (**Figure 1a**). Maps of mean fMRI activation amplitudes show that contralateral forelimb stimulation produces robust BOLD responses in cortical and subcortical somatosensory structures, with cortical signal predominating after 1400W injection (**Figure 1b**). After correction for global signal drift^10^, the difference between these datasets isolates NOSTIC-dependent components of the fMRI signal (**Figure 1c**); these responses are presumed to arise from cell populations presynaptic to the HSV-NOSTIC injection locus in POm. Such responses are correspondingly absent in fMRI data obtained from control animals treated in POm with an inert HSV-mCherry viral vector (**Figure 1d** and **Supplementary Figure 2**), ruling out sources of artifact in the measurement of projection-specific NOSTIC signals. Fore-limb stimulation response maps obtained in the control animals also resemble data obtained with NOSTIC in the presence of the 1400W inhibitor, confirming that these signals arise from intrinsic neural activation as visualized in conventional fMRI experiments.

**Figure 1.**
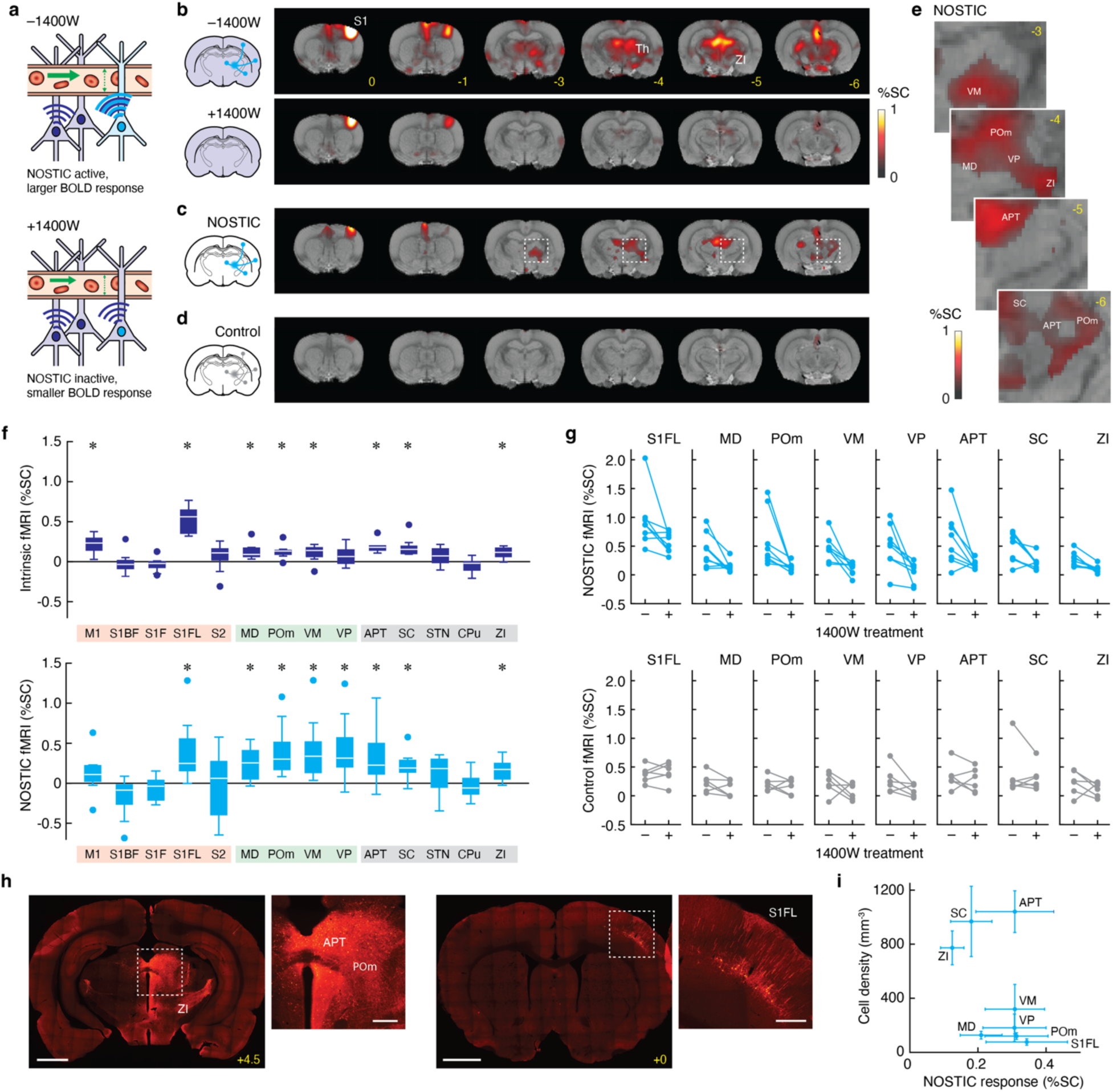
Analysis of corticothalamic circuit activity using hemodynamic imaging. **a**, Schematic of NOSTIC mechanism of action. Cells that express NOSTIC (cyan) give rise to hemodynamic contrast that can be pharmacologically modulated by 1400W. Intrinsic fMRI signals (dark blue) arise from pathways that are unaffected by the drug. **b**, Activation maps showing response to forelimb stimulation of animals treated with HSV-NOSTIC in POm. Color overlays indicate mean percent signal changes from 8 animals in the absence (top) and presence (bottom) of 1400W. Bregma coordinates in yellow. **c**, Difference map showing mean 1400W-dependent change in forelimb response for NOSTIC-expressing animals. **d**, Difference map showing 1400W-dependent change in forelimb response for control mCherry-expressing animals (*n* = 6). **e**, Closeups showing 1400W-dependent fMRI responses in thalamic subregions of NOSTIC-expressing animals. **f**, Intrinsic (top) and NOSTIC (bottom) fMRI signal amplitudes observed in cortical (pink shading), thalamic (green), and other subcortical (gray) ROIs. Box plots denote median (center line), first and third quartiles (boxes), full range (whiskers), and outliers (dots). **g**, Stimulus responses observed before and after 1400W injection in individual ROIs in NOSTIC-expressing (top) and mCherry-expressing (bottom) animals. All NOSTIC differences are significant with paired *t*-test *p* ≤ 0.05 (*n* = 8), while mCherry differences are not significant with *p* > 0.05 (*n* = 6). **h**, Representative histological data showing NOSTIC expression in key regions where hemogenetic signals are observed. Scale bars: 2 mm (full slices), 400 µm (closeups corresponding to dashed boxes). **i**, Correspondence between NOSTIC-related fluorescence and hemogenetic signals. All fMRI maps show data for voxels with significant responses or 1400W-dependent response differences with false discovery rate (FDR)-corrected *F*-test *p* ≤ 0.05 (*n* = 8).

Major features of the NOSTIC-based map of POm input include activity signatures in the S1 forelimb region (S1FL), anterior pretectal areas (APT), SC, and thalamic subdivisions including POm itself, VP, mediodorsal (MD), and ventromedial (VM) thalamic areas (**Figure 1e** and **Supplementary Figure 3**). Mean NOSTIC and intrinsic fMRI amplitudes observed in each of these regions of interest (ROIs) contralateral to the stimulus and ipsilateral to the POm viral injection site are shown in **Figure 1f**. NOSTIC and intrinsic results are correlated (*R* = 0.58, *p* = 0.027) but nevertheless vary significantly differently (two-way ANOVA, *p* = 0.023) across the 14 ROIs we examined, consistent with the expectation that NOSTIC signals arise only from a subset of the cells that give rise to intrinsic fMRI signals. Significant NOSTIC signals are observed in S1FL, MD, POm, VM, VP, APT, SC and ZI (Student’s *t*-test *p* ≤ 0.048 *n* = 8), but not in related regions such as the S1 barrel field (S1BF), S2, or primary motor cortex (M1) (*t*-test *p* ≥ 0.23). Direct comparison of stimulus response magnitudes before and after 1400W in regions with the most pronounced NOSTIC signals shows that results are consistent across animals (**Figure 1g**). Additional NOSTIC signals are apparent in regions ipsilateral to the stimulated forelimb and contralateral to the HSV-NOSTIC injection site (**Supplementary Figure 4**), indicating that transhemispheric input to POm also contributes to somatosensory response dynamics.

After neuroimaging experiments, animals were sacrificed and NOSTIC expression profiles were visualized at high resolution by fluorescence histology. As in previous tracing studies^16, 29^, results reveal dense labeling in thalamic regions in and around the HSV injection site in POm, some of which might be contaminated by spillover from the viral injection, as well as pronounced staining in distal areas including ZI, S1, and M1 (**Figure 1h**). Fluorescence levels were quantified across anatomical ROIs and compared with NOSTIC fMRI amplitudes reported above (**Figure 1i**). Correlation between these two metrics is relatively weak, however (*R* = 0.30, *p* = 0.21, *n* = 14), reflecting the fact that NOSTIC-based fMRI signals depend not only on expression of the NOSTIC probe in specific projections, but on the activity of the probe-expressing cells. A particularly high ratio of NOSTIC signal to fluorescence is observed in S1FL, consistent with the pronounced neural activity and POm input expected in this region under contralateral stimulation. In contrast, subcortical regions including APT, ZI, and SC have both very high expression levels and relatively strong NOSTIC signaling. These neuroimaging and histology results together demonstrate the ability of NOSTIC to provide strong projection-specific activity readouts when combined with a retrogradely transported viral vector in somatosensory circuitry of the rat brain.

## Source activity transforms into outputs

The data of **Figure 1** show that projection-associated neural population activity in the somatosensory system varies differently over ROIs than anatomical connectivity strengths or endogenous nonspecific fMRI responses, but it does not specify how projection-dependent signals are related to functional properties at their loci of origin. To examine these relationships, we compared tuning and temporal characteristics of NOSTIC-responsive POm inputs to activity at their source regions across the somatosensory system. We started by comparing fMRI responses to fore-limb stimuli delivered at 3 Hz, 9 Hz, and 27 Hz frequencies, which are known to produce varying responses^30–33^ (**Figure 2a**). Effects of each stimulus frequency were quantified over ROIs (**Figure 2b**). Among the regions examined, two-way ANOVA tests revealed significantly different tuning curves for NOSTIC and intrinsic responses in MD, POm, APT, VM, and VP (all *p* ≤ 0.038, *n* = 3 frequencies).

**Figure 2.**
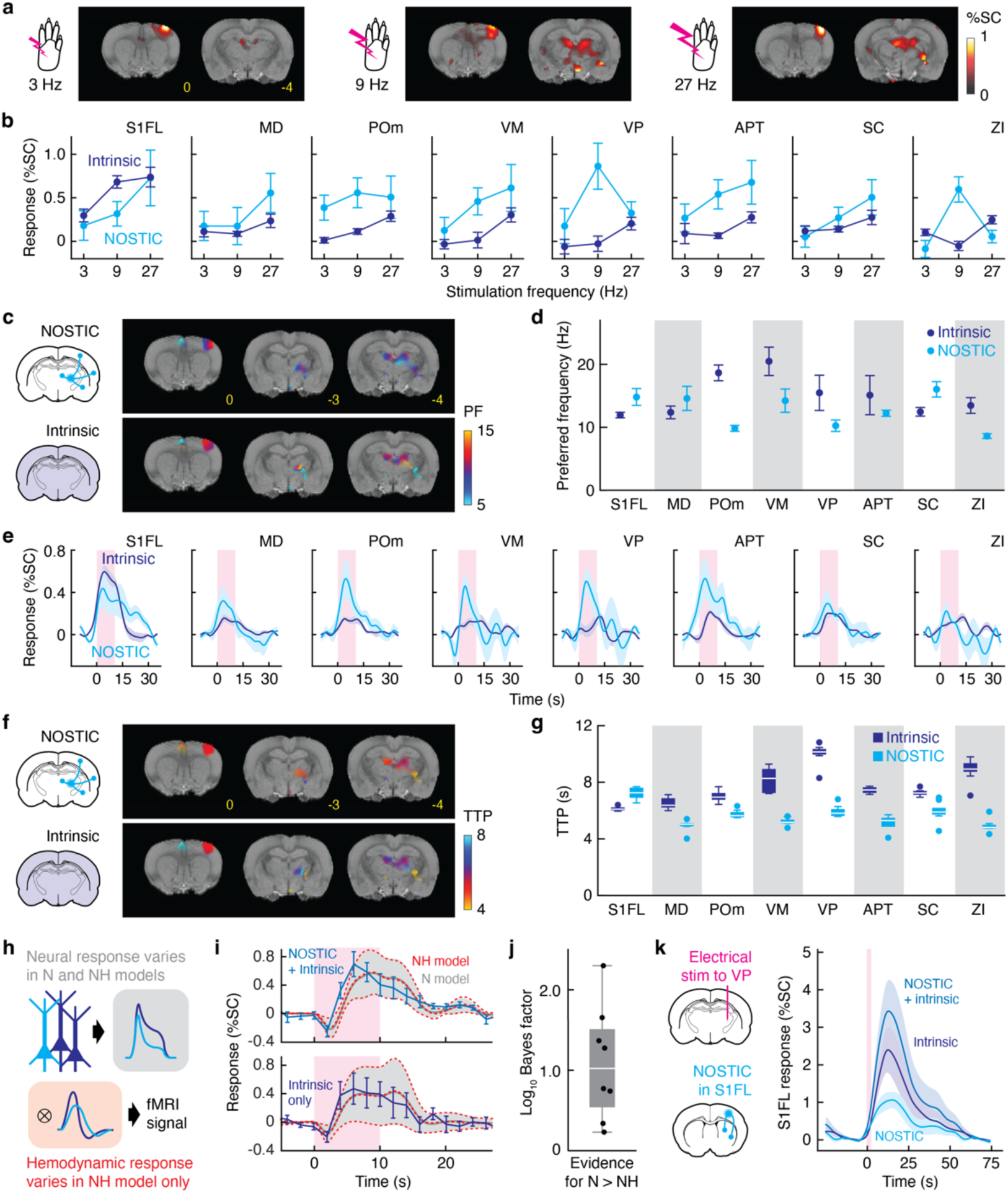
Projection-specific neural activity deviates from source regions. **a**, NOSTIC-dependent difference signal maps in response to 3 Hz (left), 9 Hz (middle), and 27 Hz (right) forelimb stimulation. **b**, Frequency-dependent NOSTIC (cyan) and intrinsic (blue) responses observed by fMRI, showing tuning in representative regions (error bars: SEM over 8 animals). **c**, Map of preferred frequency (PF) for NOSTIC (top) and intrinsic (bottom) signals in selected coronal brain sections, shown for voxels with significant NOSTIC signal (FDR-corrected *F*-test *p* ≤ 0.05). **d**, Preferred frequency for NOSTIC and intrinsic fMRI signals observed in ROIs. Dots: mean values, error bars: SEM over 8 jackknifed animals. **e**, NOSTIC and intrinsic signal time courses in response to forelimb stimulation. Stimulation period denoted by pink boxes. Shading: SEM over 8 animals. **f**, Maps showing time to peak (TTP) of NOSTIC and intrinsic fMRI responses in selected coronal brain sections. **g**, TTP values observed in regions of interest. Box plot denotes median (center line), first and third quartiles (boxes), full range (whiskers), and outliers (dots). **h**, Approach for modeling fMRI signal, in which neuronal responses from NOSTIC positive or negative components (top) are convolved with NOSTIC-independent (model N) or NOSTIC-dependent (model NH) hemodynamic responses (bottom). **i**, Response time courses showing that N (gray) and NH (red) models produce identical fits to S1 time courses observed in the absence (top) and presence (bottom) of 1400W. Error bars: SEMs over 3 animals; error margins: uncertainty from model fitting. **j**, Bayes factor describing evidence in favor of the N model over the NH model. Box plot conventions as in (g). **k**, Intrinsic + NOSTIC (–1400W), intrinsic (+1400W), and NOSTIC (difference) fMRI response time courses observed in S1 regions infected with NOSTIC-HSV under VP electrical stimulation, as diagrammed.

Maps of the stimulus tuning preferences of NOSTIC-related and endogenous fMRI signals show that these two readouts differ systematically under the conditions we applied. NOSTIC responses are biased systematically toward lower frequency stimuli (**Figure 2c**), with a mean weighted average over 1400W-dependent voxels of 8.33 ± 0.02 Hz, versus 10.71 ± 0.03 Hz for intrinsic fMRI signals—a significant difference with *t*-test *p* < 0.001 (*n* = 6.6 × 10^4^ voxels). Quantification of the weighted frequency average over ROIs shows a related trend (**Figure 2d**). Average NOSTIC signals are tuned to lower frequencies than endogenous signals in five out of eight ROIs, significantly so in ZI and POm itself (*t*-test *p* ≤ 0.036, *n* = 8 animals), although the frequency preference in SC was higher for NOSTIC than endogenous fMRI responses (*t*-test *p* = 0.033). Compared with nonspecific responses, projection-associated fMRI signals are also tuned more narrowly, a phenomenon that is significant over voxels with *F*-test *p* < 0.001 (*n* = 6.6 × 10^4^). These results thus demonstrate that neural activity at source regions is filtered to yield projection-specific signals, which themselves trend toward common characteristics as they converge onto POm.

By comparing the time courses of fMRI responses in regions with strong stimulus-evoked NOSTIC signals, we could further study how source region information is transformed into projection-associated activity. **Figure 2e** shows average forelimb stimulation response traces measured by NOSTIC and intrinsic fMRI in each relevant ROI. From such time courses, we computed mean time to peak (TTP) values for each voxel (**Figure 2f**) and brain region (**Figure 2g**). The results show that NOSTIC signals reach their maxima slightly faster on average, with mean TTP of 5.32 ± 0.01 s, versus 5.70 ± 0.02 s for endogenous fMRI responses (significant with *t*-test *p* < 0.001, *n* = 6.6 × 10^4^ voxels). At an ROI level, POm inputs from all regions except S1FL and SC displayed significantly shorter TTP values than their corresponding source regions (*t*-test *p* ≤ 0.029, *n* = 8), consistent with the sharper appearance of NOSTIC responses in the time courses of **Figure 2e**. Quantification of full width at half-maximum (FWHM) values also reveals dissociations between NOSTIC and intrinsic fMRI time courses (**Supplementary Figure 5**), though the directionality of observed changes varies among regions. Significantly shorter projection-specific response durations were seen in MD and VM (*t*-test *p* ≤ 0.005, *n* = 8), while significantly longer FWHM values were observed for NOSTIC data in S1FL and APT (*t*-test *p* ≤ 0.0002, *n* = 8). These results imply that temporal characteristics of POm inputs differ from source region activity, with faster responses generally associated with the projections.

To rule out the possibility that differences between temporal dynamics of NOSTIC-driven and endogenous BOLD responses arise simply from probe- or 1400W-dependent changes in neurovascular coupling, we used a popular modeling approach^34^ to compare characteristics of BOLD responses observed before and after 1400W (**Figure 2h**). Comparison of neural and hemodynamic parameters required to fit the two responses shows that changes in the neural terms, but not the hemodynamic variables are sufficient to explain the effects of 1400W (**Figure 2i** and **Supplementary Figure 6**), with a Bayes factor favoring the neural-only model over the neural-hemodynamic model both at the group level and in each individual animal (**Figure 2j**). This is consistent with the hypothesis that time course differences attributed to NOSTIC-labeled *vs.* unlabeled cell populations arise from activity characteristics of the labeled cells, as opposed to NOSTIC-dependent differences in the hemodynamic response.

As a further test, we performed an experimental measurement in which HSV-NOSTIC was introduced into S1FL and stimulation was applied using an intracranial electrode targeted to VP^35^, the main source of feed-forward S1 input^36^ (**Figure 2k**). In this configuration, we predicted that NOSTIC and intrinsic fMRI responses in S1FL would reflect contributions of the same neural population, and that time course differences should not arise unless NOSTIC and intrinsic hemodynamic responses diverge. Indeed, we found that fMRI signals measured in the absence and presence of 1400W correlated almost perfectly with each other (*R* = 0.98, significant with *p* < 0.001), suggesting that NOSTIC and endogenous BOLD time courses are virtually identical in these neural populations. These results further support the conclusion that the temporal distinctions documented in **Figure 2e-g** reflect differences in the activity of projection-specific versus nonspecific neural populations, as opposed to hemodynamic differences between the NOSTIC-mediated and endogenous fMRI readouts.

## Projection activity is target-specific

The fact that POm-projecting fMRI signals tend to exhibit sharper tuning and temporal characteristics than their source regions suggests that local processing functions transform regional activity into distinct output signatures from the brain areas we studied. We wondered whether such transformations lead only to a single mode of output, or whether outputs sent to different target regions also exhibit different activity profiles from one another. To address this, we expanded on the dataset of **Figure 1**, which probes inputs to POm contralateral to the stimulated forelimb, by adding data from the same eight animals on inputs to POm ipsilateral to the stimulation, as well as fMRI measurements from an additional nine rats treated with HSV-NOSTIC in S1FL, which permit activity of NOSTIC-expressing inputs to S1 to be studied.

Maps of responses to forelimb stimulation in these experiments are shown in **Figure 3a,b**, and box plots of ROI-level responses are shown in **Supplementary Figure 7**. As in the initial dataset, a combination of cortical and subcortical features are apparent in the 1400W-dependent difference data, reflecting a mix of inputs to the virally targeted structures from regions that overlap with those examined in **Figures 1-2**. POm-injected animals under ipsilateral stimulation show strong NOSTIC signals in ipsilateral POm and M1, as well as contralateral POm, MD and APT. S1-injected animals display NOSTIC responses in S1FL itself, M1, and thalamic regions. Post-mortem histological analysis confirms NOSTIC expression in these areas, though at lower levels than observed in POm-treated animals (**Supplementary Figure 8**). Probe expression is also observed in S2 ipsilateral and S1 contralateral to the HSV-NOSTIC injection sites, where 1400W-dependent signals were not observed, suggesting that these areas provide input to S1 only under conditions not examined in these experiments.

**Figure 3.**
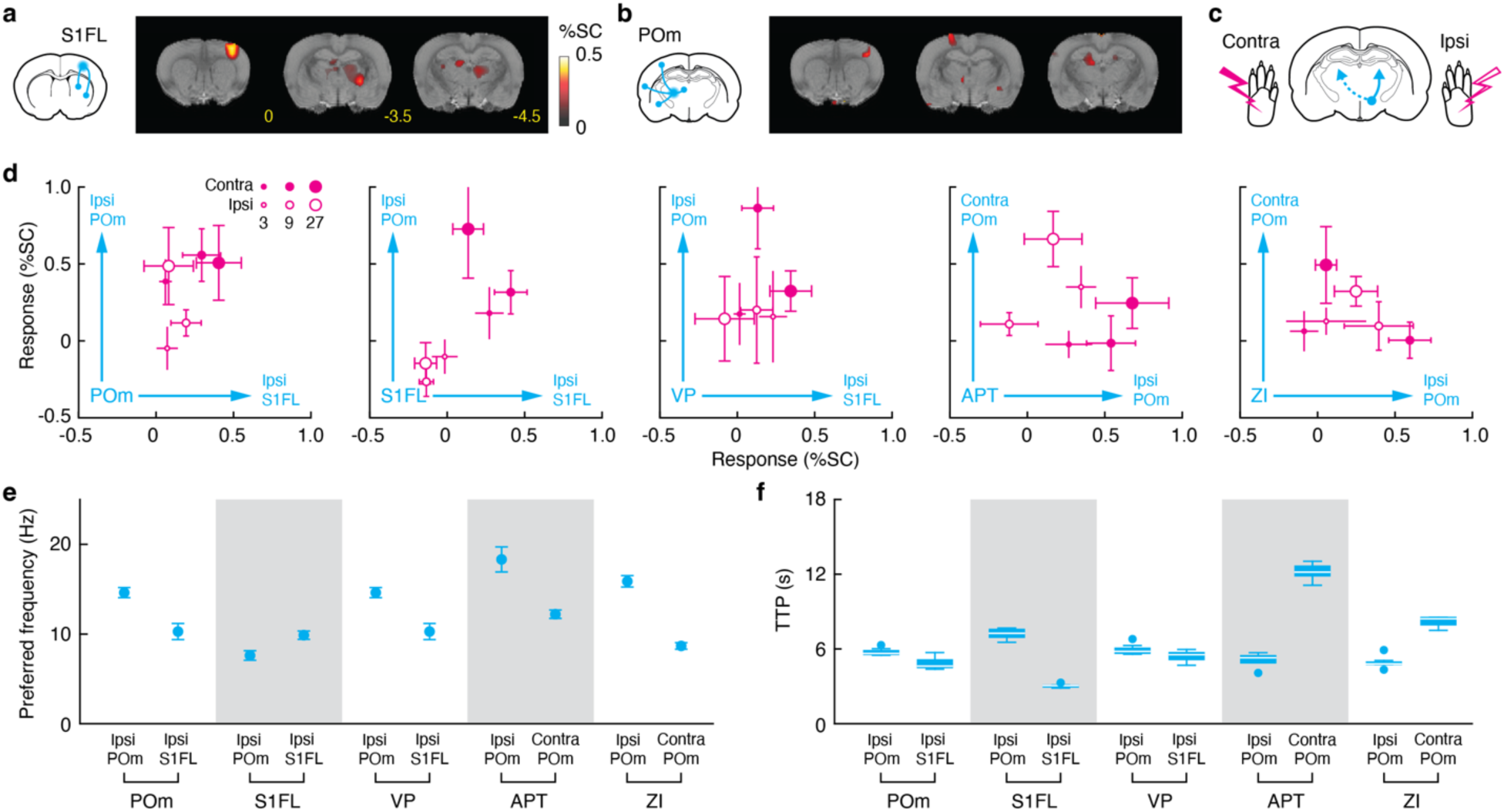
Brain regions transmit disparate information to different targets. **a**, NOSTIC fMRI measurements of input to S1FL contralateral to the stimulated forelimb in select coronal slices, with schematic at left. **b**, NOSTIC fMRI measurements of input to POm ipsilateral to the stimulated forelimb in select coronal slices, with schematic at left. **c**, Projections (blue lines) and stimuli (pink) are defined with respect to source regions, under the assumption of left-right symmetry in the somatosensory system. **d**, NOSTIC responses from projections that share the same source, measured under stimulation ipsilateral (open circles) or contralateral (filled circles) to the source region, at three stimulation frequencies (circle sizes). **e**, Preferred frequency in projections to pairs of targets (top labels) that share the same source (bottom labels). Dots: mean values, error bars: SEM over 8 or 9 jackknifed animals. **f**, Response time to peak (TTP) in projections to pairs of targets (top labels) that share the same source (bottom labels). Box plot denotes median (center line), first and third quartiles (boxes), full range (whiskers), and outliers (dots). Maps in (a) and (b) show fMRI signal changes for voxels with significant 1400W dependence (FDR-corrected *F*-test *p* ≤ 0.05, *n* = 8).

By combining measurements from inputs to contralateral POm, ipsilateral POm, and contra-lateral S1, with respect to forelimb stimulation, we could compare the activity profiles of projections from individual brain regions to targets in both hemispheres (**Figure 3c**). The dataset in particular samples projections from S1FL, POm, and VP to ipsilateral POm and S1FL, as well as presynaptic input from APT and ZI to both ipsilateral and contralateral POm. **Figure 3d** compares the mean response magnitudes of pairs of output projections that share the same source, under both contralateral and ipsilateral forelimb stimulation at three frequencies. The activity of projection pairs under these stimuli is uncorrelated in every case (|*R*| ≤ 0.66, *p* ≥ 0.15, *n* = 6 conditions). The interaction between projection target on frequency tuning is significant for outputs from S1FL (mixed two-way ANOVA *p* = 0.008, *n* = 8,9) and ZI (within-subject two-way ANOVA *p* = 0.032, *n* = 8), and the interaction of projection target with ipsi- *vs.* contralateral forelimb preference is also significant for outputs from S1FL (mixed two-way ANOVA *p* = 0.020, *n* = 8,9). Examination of the tuning and temporal characteristics of each projection pair (**Figure 3e,f**) further shows significant differences in frequency tuning for all five pairs (unpaired *t*-test *p* ≤ 1 × 10^-5^, *n* = 8-9), as well as differences in TTP for three out of five pairs (*t*-test *p* ≤ 0.016, *n* = 8-9). These results thus together show that outputs from individual somatosensory regions transmit distinct information to different postsynaptic targets in the brain.

## POm input reconfigures in different contexts

We next sought to investigate the relationships among inputs from multiple source regions that impinge onto individual target structures. By comparing projection-specific NOSTIC fMRI signals in the presence of contralateral *vs.* ipsilateral forelimb stimulation (**Figure 4a**), we readily observed that the relative amplitudes of source-specific responses differ, presumably reflecting the distinct pathways of stimulus propagation from the stimulated limb. Projection-specific response amplitudes are uncorrelated between the two stimulus conditions (*R* = 0.08, *p* = 0.79, *n* = 11 projections). Although only one individual projection showed significantly different response amplitudes under the two stimulation condition (S1FL to POm, *t*-test *p* = 0.026, *n* = 8 animals), mean projection-specific activation amplitudes were systematically greater under stimulation of the contralateral versus ipsilateral forelimb (paired *t*-test *p* = 1 × 10^-4^, *n* = 11).

**Figure 4.**
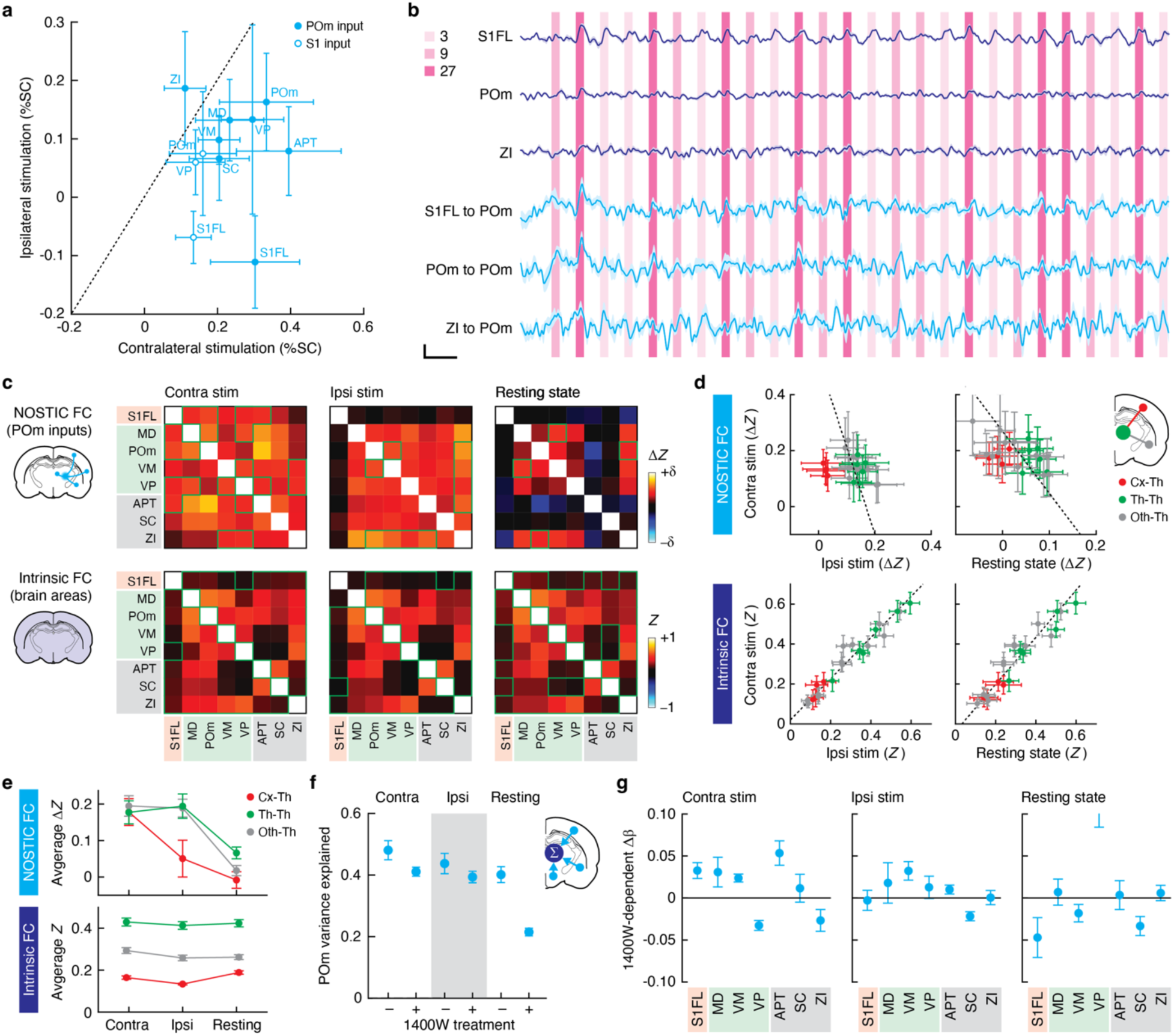
Projection activity reconfigures under different stimulus conditions. **a**, Comparison of NOSTIC-dependent signal during contralateral and ipsilateral forelimb stimulation. **b**, Mean time courses of intrinsic and hemogenetic fMRI signal for select brain regions in animals treated with HSV-NOSTIC. **c**, 1400W-dependent Δ*Z* between pairs of projections that provide input to POm (top) and corresponding *Z* values for intrinsic FC matrices (bottom) during contralateral forelimb stimulation (left), ipsilateral stimulation (middle), and resting state (right) conditions. δ = 0.4 for contra and ipsi stim, 0.1 for resting state. Green outlines denote significance of individual matrix elements with *t*-test *p* ≤ 0.05 (*n* = 8 for contra and ipsi stim, *n* = 9 for resting). **d**, Comparison Δ*Z* values for S1 inputs (top) and *Z* values for intrinsic FC measures (bottom) among stimulation conditions. Color coding distinguishes cortico-thalamic FC (red), thalamothalamic FC (green), and FC between thalamic and other subcortical regions (gray). Dots: mean, error bars: SEM over 8 or 9 animals. **e**, Average Δ*Z* and *Z* values for different classes of connections under different stimulus conditions, following the color coding of (d). Error bars: SEM over 4-12 matrix elements. **f**, Fraction of variance explained by a GLM that estimates POm fMRI signal in terms of a sum of inputs from source regions where NOSTIC signals are observed (schematic at right), either in the absence or presence of 1400W. Dots: means, error bars: SEM over 8 or 9 jackknifed animals. GLMs in the absence of NOSTIC inhibition perform significantly better in contra and resting conditions (*t*-test *p* ≤ 0.034). **g**, 1400W-dependent difference in GLM weights for POm inputs during contralateral forelimb stimulation (left), ipsilateral stimulation (middle), and rest (right). Dots: means, error bars: SEM over 8 or 9 jackknifed animals.

The relative amplitudes of neural activity in specific projections depend on how the neural system is excited, but we hypothesized that the correlational structure of inputs to individual structures might be more conserved, due to the relatively fixed nature of physical connectivity among brain regions on the time scale of our measurements. We also anticipated that sensory stimulation of any kind might drive highly correlated input to each brain region, even under different sources of sensory drive. Indeed, conservation of FC across stimulated and stimulus-free (resting) states has been reported in a number of human studies^37–39^. In order to examine whether activity in NOSTIC-labeled projections would exhibit invariant correlational patterns, we evaluated interregional correlation among extended time courses of fMRI signals such as those represented in **Figure 4b**. By calculating the 1400W-dependent differences in *Z*-transformed correlation coefficients, we could determine NOSTIC-dependent contributions to the interregional correlations. The resulting Δ*Z* values from paired regions represent correlated input from NOSTIC-labeled projections, as well as correspondences between NOSTIC signal in one region and endogenous signal in the other region. We conducted this analysis only on rats treated with HSV-NOSTIC in POm, because of the superior probe expression and rich set of projections labeled in these animals.

NOSTIC FC matrices containing Δ*Z* values among POm inputs observed under contralateral stimulation, ipsilateral stimulation, and resting state conditions are shown in **Figure 4c**, along with intrinsic FC matrices computed from endogenous regional signals observed in the same animals after addition of 1400W; **Supplementary Figure 9** shows analogous pre-1400W data. The NOSTIC FC matrices reveal strikingly different patterns among the three conditions we assessed, whereas the endogenous FC matrices are virtually unchanged across conditions, as are FC matrices from control HSV-mCherry-treated animals (**Supplementary Figure 10**). NOSTIC-dependent Δ*Z* values for POm inputs observed under contralateral stimulation are anticorrelated with those observed during rest (*R* = –0.52, *p* = 0.012, *n* = 22 projection pairs) or ipsilateral stimulation (*R* = –0.26, *p* = 0.18), reflecting a restructuring of information flow to POm under the contrasting conditions (**Figure 4d**). In contrast, *Z* values for intrinsic FC are highly positively correlated among the three conditions (*R* = 0.95-0.97, *p* ≤ 0.001, *n* = 22). Among NOSTIC-labeled POm inputs, the FC between cortical and thalamic projections is particular dependent on conditions (**Figure 4e**), with Δ*Z* values that were significantly stronger under contralateral stimulation than ipsilateral stimulation or rest (paired *t*-test *p* ≤ 0.02, *n* = 4). FC among thalamic projections or between thalamic and other subcortical projections is strong under stimulation, but significantly weaker in the resting state (*t*-test *p* ≤ 0.002, *n* = 6-12). Again, these context-dependent effects are not observed in the intrinsic FC data, which show relatively constant *Z* values across all conditions. Our results therefore emphasize that POm receives disparate combinations of input from different brain sources, and that information flow patterns reorganize in stimulus-dependent fashion.

As a complement to correlational analysis, we used general linear modeling (GLM) analysis^40^ to estimate the relative contributions of different input sources to population activity in POm. To do this, mean time courses of POm fMRI signal were modeled in terms of time courses from each brain region where significant NOSTIC signals were observed in **Figure 1**. GLMs computed using this approach fit data obtained in the absence of 1400W better than in the presence of 1400W, indicating that NOSTIC signals in the source regions can help predict the fMRI time courses in POm (**Figure 4f**). This was particularly true in the resting state, where the inhibition of NOSTIC decreased the fMRI variance explained in POm by close to 50%, a significant effect with unpaired *t*-test *p* = 9 × 10^-5^ (*n* = 8 animals). To estimate the magnitudes of contributions from specific projections, 1400W-dependent differences in regression coefficients were computed for each model. Values for contralateral stimulation, ipsilateral stimulation, and rest differed significantly from one another, with two-way ANOVA *p* < 0.001 over *n* = 7 ROIs. NOSTIC regression coefficients were uncorrelated across the conditions (|*R*| ≤ 0.59, *p* ≥ 0.16, *n* = 7), whereas an equivalent analysis on intrinsic fMRI signals (**Supplementary Figure 11**) results in GLM coefficients that are highly correlated between contralateral and ipsilateral stimulation conditions (*R* = 0.97, *p* = 0.0003). These results therefore further support the finding that information flow patterns in the somatosensory system undergo stimulus-dependent reorganization, and that these changes are not detectable from intrinsic fMRI data alone.

## Network E/I balance changes dynamically

Despite the anatomical and functional differences among projections we studied, it is striking that none of the NOSTIC fMRI amplitudes or the correlations involving them are significantly negative in any of the conditions we addressed (*t*-test *p* > 0.35, *n* = 8-9 animals). This suggests that neural populations with contrasting functional roles remain balanced with one another at a mesoscopic level throughout the network, in analogy to the well known phenomenon of excitation/inhibition (E/I) balance in cortical microcircuitry^41, 42^. To explore this idea, we quantified the fraction of inhibitory cells in each NOSTIC-labeled neural population (**Figure 5a**). Inhibitory POm-projecting cells were identified as those that stain for the mCherry tag fused to NOSTIC and that express mRNA for the vesicular γ-aminobutyric acid transporter (VGAT). Consistent with literature^16^, results of this analysis reveal that projections to POm from MD, VM, APT, SC, and ZI are more inhibitory than excitatory, with input from ZI to POm in particular composed more than 75% of VGAT-positive cells (**Figure 5b**). Given that activity of both excitatory and inhibitory POm inputs remains positively correlated across conditions (**Figure 4c**), these data indicate a phenomenon of network-wide E/I balance among projections to POm.

**Figure 5.**
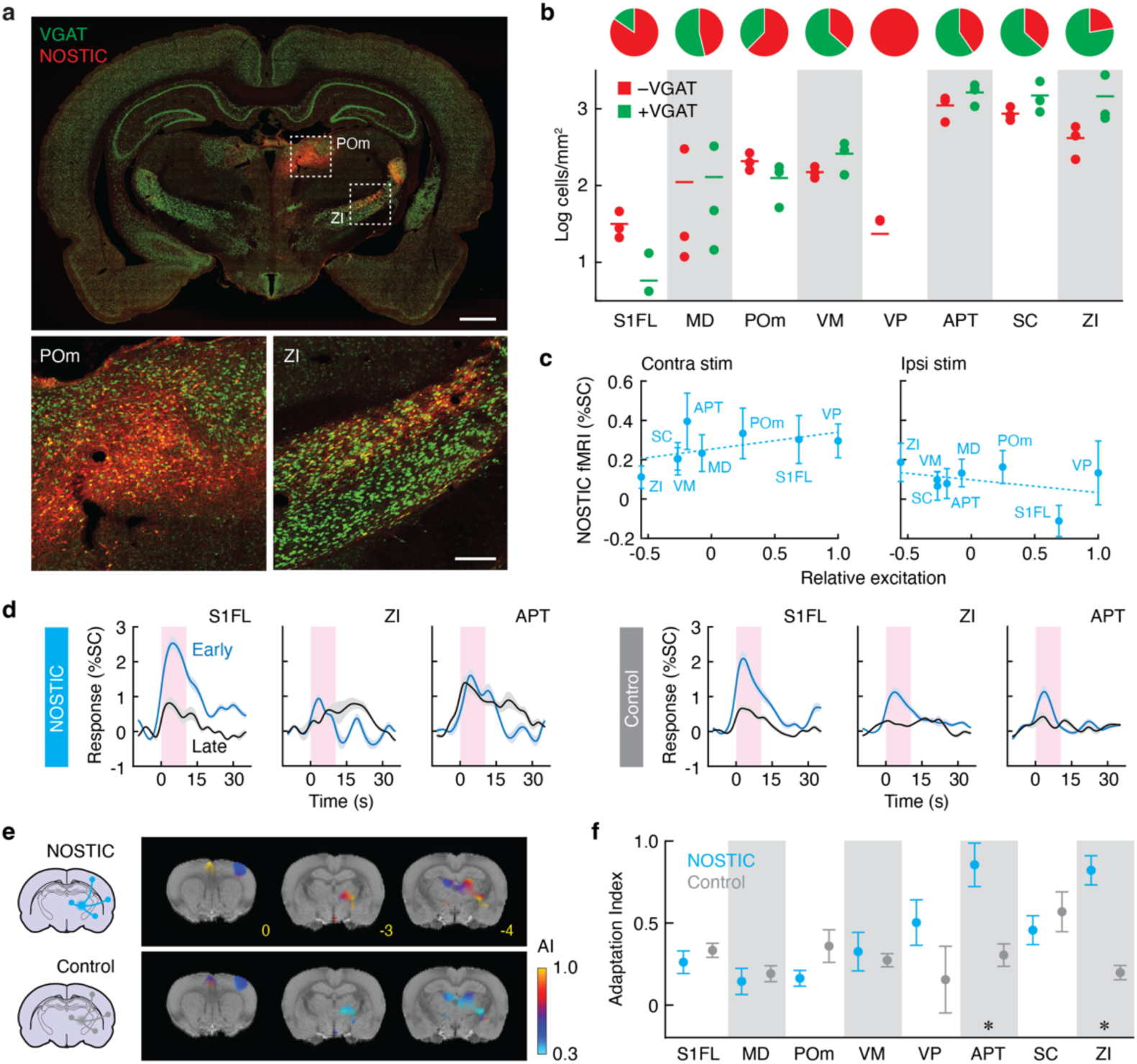
Network-wide E/I balance contributes to response tuning and adaptation. **a**, Visualization of NOSTIC expression with respect to a marker of inhibitory neurons (VGAT) in representative postmortem brain slices. Close-ups (bottom) correspond to dashed regions at top. Scale bars: 1 mm (top), 200 µm (bottom). **b**, Quantification of labeled cell densities in brain regions with NOSTIC responses. Data points indicate densities of NOSTIC-positive and VGAT-negative cells (red) and NOSTIC-positive VGAT-positive cells (green) from three independently processed animals. **c**, Comparison of relative excitation estimated from staining results with NOSTIC fMRI signals observed under forelimb stimulation contralateral (left) and ipsilateral (right) to NOSTIC injection and measurement regions (labeled). Dashed lines indicate positive (left) *vs.* negative (right) trends in the relationships between relative excitation and NOSTIC signal. **d**, Early (blue) and late (black) responses to repeated 27 Hz stimulation in three regions of interest (labeled), for animals treated with HSV-NOSTIC (left) or with the HSV-mCherry control virus (right). Curves reflect the mean response to the first or last two stimuli in a series of ten repetitions, averaged over 8 (NOSTIC) or 6 (control) rats; shading denotes SEM. Pink boxes denote the stimulation period. **e**, Voxel level maps of adaptation indices (AI) for animals treated with POm injections of HSV-NOSTIC (top) or control HSV-mCherry (bottom). Color coding indicates AI for voxels showing significant NOSTIC signal (FDR-corrected *F*-test *p* ≤ 0.05). **f**, Adaptation indices determined for ROIs indicated in NOSTIC (cyan) or control (gray) animals. Means (dots) and SEM values (error bars) were computed from jackknifed datasets.

Dynamic changes in E/I balance are known to regulate activity levels in neocortical circuitry^41, 42^. To examine how this principle contributes to stimulus processing in at a systems level, we estimated the relative excitation contributed by each input to POm as (*E* – *I*)/(*E* + *I*), where *E* and *I* are the numbers of VGAT-negative and VGAT-positive NOSTIC-labeled cells, respectively. Plotting relative excitation against NOSTIC-dependent signal changes observed under contralateral versus ipsilateral forelimb stimulation shows how the balance of excitatory to inhibitory inputs shifts between these conditions, favoring inhibitory input to POm when the disfavored limb is stimulated (**Figure 5c**). Slopes of the fitted trend lines are opposite under the two conditions, with the most inhibitory POm input (from ZI) more activated under ipsilateral stimulation while the most excitatory inputs (S1FL and VP) are more activated under contralateral stimulation. This suggests a role for changing E/I balance in tuning for the preferred forelimb.

Shifts in E/I balance also accompany sensory adaptation^43–45^, but few studies have examined this phenomenon at a multiregional network level. We looked for evidence of system-wide shifts in E/I balance during adaptation by comparing fMRI responses to contralateral forelimb stimulation at early versus late time intervals over the course of repeated 27-Hz stimuli. To estimate NOSTIC-dependent components without confounds due to the time between measurements before and after 1400W administration, we compared adaptation over equivalent pre-1400W time intervals between test rats that received POm injections of HSV-NOSTIC and control animals that were treated instead with the inert HSV-mCherry vector.

Figure 5d shows mean responses in the first versus last two cycles in a 10-cycle series of 27-Hz stimuli lasting 16 min for three representative ROIs where viral labeling is observed, showing pronounced adaptation in multiple regions for the control animals, but much less adaptation in some ROIs for the NOSTIC-treated animals. We used the response amplitudes from such traces to compute an adaptation index as the ratio of late to early response magnitudes, reported over voxels in Figure 5e and over regions in Figure 5f. This analysis shows that ZI and APT inputs to POm both show significantly less adaptation in NOSTIC than in mCherry animals (unpaired *t*-test *p* ≤ 0.033, *n* = 8 NOSTIC, 6 mCherry rats), while other projections show strong adaptation (values below ∼0.5 for both test and control). Interestingly, the two regions that show least adaptation are also among the regions that provide the highest proportion of inhibitory input to POm (Figure 5b). This shows that adaptation to forelimb stimulation is accompanied by a substantial increase in the relative inhibitory input to POm, reflecting a network-level shift in E/I balance throughout the somatosensory system.

## Discussion

In a brain-wide fMRI study of projection-specific function in the rodent somatosensory system, we discovered three principles by which mesoscale regional activity is transformed into output and recombined into input at cortical and subcortical sites. First, population activity profiles of outputs from individual brain regions differ from their sources in tuning and timing, displaying target-dependent characteristics. Second, the pattern of inputs from multiple source regions onto an individual target area alters dramatically as a function of stimulation or rest, without corresponding changes in conventionally defined functional connectivity. And third, excitatory and inhibitory projections are positively correlated in all conditions we studied, demonstrating the phenomenon of system-wide E/I balance, but this balance shifts toward inhibition over the course of repeated stimulation or in the presence of non-preferred stimuli. These results show how multiregional interactions in the brain contribute to fundamental phenomena such as stimulus tuning and adaptation. They also demonstrate the utility of measuring neural information flow directly, rather than inferring input/output relationships based on connectivity and regional activity alone.

The analyses reported here made particular use of the genetically encoded neural activity reporter for fMRI called NOSTIC^10^. By applying the probe in a retrogradely transported viral vector^13^, we could perform simultaneous activity measurements from distributed brain regions pre-synaptic to an injection site, under both stimulated and resting state conditions. This approach differs substantially from network-level fMRI studies that combine circuit-specific activity manipulations with imaging^46–48^, where perturbations but not readouts are well defined. It also offers more comprehensive spatial coverage than invasive techniques such as electrophysiology with projection-specific optogenetic tagging^49, 50^, facilitating the observation of brain-wide phenomena such as the correlated patterns of information flow we document in Figure 4. At the same time, limitations of the technique suggest caveats to the current study. First, because the NOSTIC probe is detected by means of hemodynamic readouts, it is subject to nonidealities arising from properties of the BOLD contrast^51–53^; here we controlled for some of these by comparing NOSTIC results with those obtained with an mCherry-encoding vector, by correcting for physiological drift using a normalization approach, and by showing experimentally that the NOSTIC hemodynamic response closely parallels the endogenous fMRI response. Second, although the comparison of histology with NOSTIC-dependent signals suggests that even sparse neural populations can be recorded with the probe, it is not yet known whether resulting fMRI signals may be biased toward specific cell types, analogous to how electrophysiology recordings tend to be biased toward pyramidal neurons in cortex^54, 55^. Third, as with all virus-dependent methods, there is the possibility that results may be skewed by nonideal behavior of the vector or its application^55^; for instance, sites close to the injection targets in POm or S1FL may be contaminated by direct spreading from the injections themselves, as well as partial volume effects arising from proximity, though major conclusions of this study would be unlikely to be affected by this.

Our experiments focused most prominently on projections presynaptic to POm, a high order thalamic region whose function is incompletely understood but thought to be involved in sensory perception, plasticity, and sensorimotor integration^56^. Although our experiments were performed in anesthetized animals, our system-wide measurements provide notable evidence of convergence of inputs onto POm that could aid in hypothesized roles in stimulus discrimination and choice on time scales of seconds to minutes^57–60^. We discovered network-wide E/I balance as a notable feature of POm inputs and we identified output from ZI as an inhibitory projection that provides particularly strong POm input under non-preferred ipsilateral forelimb stimulation and that fails to adapt over repeated stimulation. The behavioral importance of afferents from ZI and other regions depends on the subsequent ability of POm to relay responses to other cortical and subcortical areas. Consistent with such a capability, we found that NOSTIC-labeled recurrent POm neurons display strong adaptation and stimulus selectivity consistent with input to POm from other areas; moreover, NOSTIC signals from S1-projecting POm output neurons display response characteristics similar to POm inputs, with comparatively low-frequency stimulus tuning and fast responses. But more complete exploration of POm output and its ability to influence circuitry beyond S1FL would benefit from the use of anterograde NOSTIC vectors that were not available for the current study. Characterization of inputs and outputs from additional somatosensory processing regions would also be possible in future experiments.

Although our results were obtained in rats, they are broadly relevant to analyses in human subjects. Our finding that projection-specific activity frequently deviates from fMRI signals in the source ROIs challenges popular models of multiregional neural integration^61^ in which the effect of each source region on its projection targets is assumed to track average population activity in the source region. Our results instead argue for modeling approaches that treat each region’s outputs as separate neural subpopulations with distinct activity profiles—perhaps by preferentially parametrizing the edges rather than the nodes of traditional graph models^62^. The resulting additional complexity could be mitigated by applying perturbations that help disambiguate network relationships, or by performing parallel animal studies using NOSTIC fMRI or related techniques. A second point of relevance follows from the surprising finding that NOSTIC-dependent FC in thalamocortical circuitry shifts under different stimulation contexts while intrinsic fMRI-related FC remains consistent. This implies that direct connectivity between pairs of regions probed in our study plays a relatively minor part in determining the overall correlational structure that conventional FC reveals. In some cases, FC measures from human neuroimaging studies may similarly embody a superposition of causal contributions, requiring cautious interpretation. Finally, our results reflect on efforts to infer the effects of inhibition in human neuroimaging studies. Human studies often consider inhibition as a local effect, reflected in cortical regions by average GABA concentrations or GABA receptor levels^63–65^, but our findings suggest that distinct long-range inhibitory projections in subcortical circuitry can be quite important for network-wide effects such as stimulus selectivity and adaptation. Further studies may characterize neurochemically distinct aspects of brain-wide activity more completely, providing deeper insights into how macroscopic aspects of brain function arise from combinations of underlying cellular-level processes.

## Abbreviations

APT: Anterior Pretectal Nucleus
BOLD: Blood Oxygen Level-Dependent
E/I: Excitation/Inhibition
FWHM: Full Width at Half-Maximum
FC: Functional Connectivity
fMRI: Functional Magnetic Resonance Imaging
HSV: Herpes Simplex Virus
MRI: Magnetic Reso-nance Imaging
M1: Primary Motor Cortex
MD: Mediodorsal Thalamus
NOSTIC: Nitric Oxide Synthase for Targeting Image Contrast
Pom: Posteriomedial Thalamus
ROI: Region of Interest
S1BF: Primary Somatosensory Cortex Barrel Field
S1FL: Primary Somatosensory Cortex Forelimb Region
S2: Secondary Somatosensory Cortex
SC: Superior Colliculus
TTP: Time To Peak
VM: Ventromedial Thalamus
VP: Ventroposterior Thalamus
ZI: Zona Incerta

## Acknowledgements

This research was funded by awards R01 NS121073, R01 DA062195, UG3 MH126868, and U01 EB031641 from the US National Institutes of Health, as well as a grant from the Simons Center for the Social Brain at MIT, all to AJ. YJ and SB were funded by J. Douglas Tan Fellowships and YK was funded by a Y. Eva Tan Fellowship, all from the McGovern Institute for Brain Research at MIT. JM was funded by an fellowship from the Integrative Computational Neuroscience Center at MIT and WT was funded by a fellowship from the Simons Center for the Social Brain. The authors are grateful to Ke Chen and Fan Wang for assistance with *in situ* RNA analysis and thank Mriganka Sur and Robert Desimone for comments on the manuscript.

## Author contributions

The first two co-authors contributed equally. YJ and YK performed the fMRI experiments with help from SB, SKC, and WT. YJ performed the fMRI analysis with help from AD. YK performed histology with help from BS and JW. JW performed histology data analysis. JM performed BOLD response modeling. YJ and AJ wrote the paper with help from the other authors.

## Author information

The authors declare no competing interests. Correspondence and requests for materials should be addressed to AJ.

## METHODS

### Animal subjects

All animal procedures were performed in strict compliance with US Federal guidelines, with oversight by the MIT Committee on Animal Care under approved protocol number 0721-059-24. Male and female Sprague-Dawley rats (250-350g) were acquired from Charles River Laboratories (Wilmington, MA).

### Intracranial viral injection

Animals were prepared for cranial surgery to expose the desired viral injection site at the POm or S1. Each animal was anesthetized with isoflurane (2% for induction, 1.5% for maintenance) and positioned in a stereotaxic instrument (Kopf Instruments, Tujunga, CA) with a water heating pad (Braintree Scientific, Braintree, MA) to keep the body temperature at 37 °C. The eyes of each animal were protected with Paralube Vet Ointment (Dechra Veterinary Products, Overland Park, KS) to prevent drying. The head was then shaved and cleaned with alcohol and povidone-iodine pads. To expose the target virus injection sites, the skin over the skull was retracted, and the skull was thoroughly cleaned using sterile surgical equipment. Holes were drilled into the skull above the target sites, which were located in the POm (bregma coordinates: AP –3.3 mm, ML ± 2 mm, DV –5.6 mm) or S1 (AP ± 0.5 mm; ML ± 3.3 mm, DV –1.8 mm).

HSV-NOSTIC and HSV-mCherry vectors were adapted from published sequences^10^ and packaged by viral cores at the Massachusetts General Hospital and University of North Carolina. Each viral vector at a titer of 1.5-2.5 × 10^9^ infectious units per mL was preloaded into a pulled glass pipette. Each vector was injected in a total volume of 2 µL for POm, and 4 µL for S1. The glass pipette was slowly removed 10 min after the viral injection. Bone wax was applied to each injection site using a sterile cotton applicator tip. Skin incisions were closed using surgical suture, and 2% lidocaine gel was applied over the wound areas. Isoflurane was then discontinued, and each rat was removed from the stereotaxic frame and placed in a warmed cage on a heating pad to recover for 45 min. Slow-release buprenorphine (0.3 mg/kg) was administered subcutaneously to minimize pain and discomfort.

### Preparation of animals for functional imaging

Immediately prior to imaging experiments, rats were anesthetized using 3% isoflurane and maintained at 2% isoflurane during preparation. Animals were intubated and ventilated (60 strokes/min, 9.5 mL stroke volume) and a tail vein catheter was established for drug delivery. Animals were then placed onto an custom made imaging cradle and secured in place using custom 3D printed earbars. 1 mg/kg pancuronium and 0.1mg/kg dexmedetomidine was administered, followed by continuous delivery of 2 mg/kg/h pancuronium and 0.02mg/kg dexmedetomidine thereafter. Respiration, heart rate, and blood oxygen saturation were monitored, and temperature was maintained with a circulating warm water for the remainder of the procedure. Isoflurane was then stopped and the animal was given at least 30 min to recover before imaging.

### Forelimb stimulation experiments

Bipolar electrical stimulation was delivered to the forelimb as sensory input using a constant-current stimulus isolator (World Precision Instruments, Sarasota, FL). Needle electrodes were inserted into left and right forelimbs to administer constant-current stimuli in pulses of 1 ms, at an intensity of 2 mA. To evaluate the tuning properties of the sensory system, stimulation was applied at three frequencies: 3, 9, and 27 Hz. Stimuli were presented in pseudorandomized 10-second blocks, with each block followed by a 20-second rest interval. Identical pseudorandomized stimulation sequences were applied in each scan series. A total of 10 stimulation blocks were delivered for each frequency during the scan series. Resting state fMRI data were collected without stimulation presentation. Each scan session consisted of a 15-min resting state series, followed by two series of 15-min each, during which stimuli were delivered to the left and then right forelimb; then there was a 30-min gap during which the NOSTIC inhibitor 1400W (25 mg/kg, IV bolus) was administered, and then a further 15-min resting state scan series followed by two additional 15-min scan series with left and right forelimb stimulation.

### Intracranial stimulation experiments

Three rats used solely for the experiment of Figure 2h were injected with 4 µL of HSV-NOSTIC in S1. Four weeks after viral injection, rats underwent a second surgery to implant electrodes for stimulation of VP. The scalp was retracted and target skull site exposed as for viral surgery. After creating a craniotomy over the right VP (AP +3 mm, ±3 mm ML, and DV –6 mm), a custom-fabricated silver bipolar electrode was carefully implanted into the targeted region. C&B Metabond dental cement (Parkell, Inc., Edgewood, NY) was applied to the entire exposed skull surface to secure the electrode in place. Immediately following surgery, animals were prepared for functional imaging. Bipolar electrical stimulation was delivered via the VP electrode using a constant-current stimulus isolator (World Precision Instruments). Stimulation was applied as bursts of 0.1 µA, 1 ms pulses delivered at 150 Hz for 500 ms, each followed by a 29.5-s rest interval. A total of 30 stimulation bursts were delivered in two 16-min experimental sessions, conducted before and after 1400W injection.

### Magnetic resonance imaging

MRI data were acquired using a 7 T Biospec MRI scanner (Bruker, Ettlingen, Germany) equipped with custom surface coil made in house. A rapid acquisition with refocused echoes (RARE) pulse sequence was employed to obtain *T*_2_-weighted anatomical images with the following parameters: number of averages (NA) = 12, matrix size = 256 × 256, field of view (FOV) = 2.56 cm × 2.56 cm, slice thickness = 1 mm, repetition time (*TR*) = 2500 ms, effective echo time (*TE*) = 44 ms, and RARE factor = 8. Functional imaging was typically performed using an echo planar imaging (EPI) pulse sequence with a matrix size of 64 × 64, FOV = 2.56 cm × 2.56 cm, slice thickness = 1 mm, *TR* = 2500 ms, and effective *TE* = 16 ms.

### Image data analysis

Images were reconstructed using Paravision 6 software (Bruker) and subsequently analyzed with AFNI^66^ and in-house Python scripts. Preprocessing of EPI data involved several steps: signal outside the brain was removed using AFNI’s *3dSkullStrip* function, motion correction was performed with AFNI’s *3dvolreg* function and volumes exhibiting displacements greater than half the voxel size were excluded from further analysis. Alignment to anatomical scans was achieved using AFNI’s *3dAllineate* function. Spatial smoothing was applied to a full width at half-maximum (FWHM) of 1 mm using AFNI’s *3dBlurToFWHM* function, and three-point median temporal smoothing was performed using AFNI’s *3dTsmooth* , and voxel intensity was normalized to its mean signal.

For forelimb stimulation datasets, voxel-wise linear regression was conducted to generate statistical maps using AFNI’s 3dDeconvolve function. To account for systematic changes in hemodynamic amplitudes between pre- and post-1400W conditions, fMRI responses obtained after 1400W treatment were renormalized to ensure that the forelimb response amplitude in the S1FL region during contralateral stimulation matched the corresponding amplitude observed prior to 1400W treatment. Difference maps were then generated by subtracting the mean response amplitude map obtained after 1400W treatment from the renormalized mean response amplitude map from the same animals before 1400W treatment.

Activation maps and functional connectivity (FC) data were computed for each individual animal. Activation maps from scan series with stimulation were thresholded with a false discovery rate (FDR)-corrected *F*-test value of *p* ≤ 0.05 for significant contribution of the relevant stimulus regressor in a full model consisting of six motion regressors and one stimulus regressor for overall activation maps or three frequency-dependent stimulus regressors for frequency-specific maps. Maps of response parameters such as preferred frequency, time to peak (TTP), full width at half-maximum (FWHM), and adaptation index (AI) are presented for voxels that pass the *F*-test *p* ≤ 0.05 threshold for significant NOSTIC signal in the single-stimulus regression model, as in Figure 1c. Color overlays in each fMRI amplitude or parameter map fade from full opacity to full transparency across the bottom half of the intensity scale defined by response amplitude.

For resting-state functional connectivity (FC) analysis, motion and global signal were first regressed out of each dataset using AFNI’s *3dDeconvolve* function. Region of interest (ROI) time courses were extracted with AFNI’s *3dROIstats* function, with ROIs defined based on a standard brain atlas (https://www.nitrc.org/projects/whs-sd-atlas/) and diagrammed in **Supplementary Figure 2**. Correlation coefficients between time courses were computed using Pearson’s correlation formula. To generate FC matrices for ROIs in which NOSTIC signal was observed, these were then converted to *Z* values using the Fisher *Z* transformation. Δ*Z* values reflecting NOSTIC contributions to FC were computed from the difference of *Z* values obtained before and after 1400W treatment. Significance of *Z* or Δ*Z* values was evaluated using paired Student’s *t*-tests over animals, with uncorrected threshold of *p* = 0.05.

For all forelimb stimulation experiments, animals included analyses and figure preparation were those that showed at least 0.3% S1FL response amplitudes in all four stimulation series (left and right, both before and after 1400W injection); this yielded 8 male and 9 female rats in total. For each of the included animals, stimulus response magnitudes for each voxel were defined by voxel-wise regression coefficients for the stimulus regressors; ROI-level response magnitudes represent average response amplitude from all voxels in the atlas-defined ROI. Intrinsic fMRI response magnitudes were defined by voxel- or ROI-level regression coefficients observed in the post-1400W series only. Difference magnitudes reflecting NOSTIC contributions were defined by differences between coefficients from pre-1400W and post-1400W sessions for individual voxels or ROIs. For analyses of Figure 5, estimated drive values were estimated by multiplying ROI-level difference magnitudes by (*E* – *I*)/(*E* + *I*), where *E* and *I* are the fractions of excitatory and inhibitory neurons associated with the projection, according to histological analysis.

Stimulus tuning maps were generated by associating intrinsic or NOSTIC fMRI response magnitudes at each frequency (3, 9, and 27 Hz) with a single color channel and presented directly. In addition, a preferred frequency value was calculated for each brain region as a logarithmically weighted average given as exp{ Σ*_i_* (*R_i_* log *f_i_*) / Σ*_i_ R_i_* }, where *i* indexes the three frequencies of stimulation *f_i_* and *R_i_* values are the corresponding observed response amplitudes. Regional response time courses were obtained by averaging signal across all voxels within each ROI; graphs depict averages and standard error of the mean (SEM) over animals. For temporal analysis, the average response time courses were fitted with a gamma function, and FWHM and TTP values for each response were derived from the fitted function for each voxel or ROI in each animal. All box plots of values indicate median (center line), first and third quartile boundaries (box limits), and full ranges excluding outliers (whiskers). Outliers are defined as data points more than 1.5 times the first-to-third interquartile range below or above the first and third quartile boundaries, respectively, and are depicted individually.

To perform general linear modeling (GLM) analysis of contributions to POm signal, the target time course in POm (**y**) was modeled as a linear combination of the time courses from input ROIs, according to the formula **y** = **βX** + **ε**, where **X** denotes a matrix of ROI-averaged time courses from each input region, **β** is a vector of regression coefficients (one value per ROI), and **ε** is a residual error vector. Least-squares optimized regression coefficients were computed by matrix inversion using the *statsmodels* library in Python. Group data were analyzed by concatenating ROI time courses from all animals with the leave-one-out method (jackknifing), and GLM parameters were calculated from each jackknifed dataset before and after 1400W administration. For each ROI, 1400W-dependent changes in GLM weights were calculated by subtracting post-1400W weights from pre-1400W weights. SEM values for regression parameters in this analysis were computed from the jackknifed datasets as the square root of 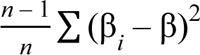, where β*_i_* is the parameter estimate obtained from the *i*th jackknife sample, β is the mean over all samples, and *n* is the total number of animals contributing to the dataset.

To analyze adaptation effects, average ROI-level time courses for pre-1400W responses to the first two and last two 27-Hz stimuli (out of 10) were generated for animals treated with NOSTIC or mCherry viruses. Response time courses were obtained from jackknifed datasets consisting of leave-one-out averages, and adaptation indices for voxels and ROIs were defined as average of the first two 27-Hz responses divided by average of the last two responses.

### Hemodynamic parameter modeling

As a basis for hemodynamic parameter modeling in Figure 2, the average time course in both S1 regions was extracted for each animal included in the analyses of Figure 1. We then fit the responses using two different variants of a BOLD response model^34^ in which an excitatory-inhibitory neural population pair was coupled to a viscoelastic balloon model^67, 68^. In the first model variant, which we term the neural model (N), differences between pre- and post-1400W sessions were assumed to arise solely from changes in the local excitatory-inhibitory balance. In the second model variant, termed the neural-hemodynamic model (NH), we additionally allowed hemodynamic parameters of the balloon model to vary across sessions. All parameters were equipped with uninformative Gaussian priors and we specified a Gaussian likelihood over the predicted time courses. Values for prior means, variances, and the observation noise covariance were extracted from the literature^34^. Parameter distributions and model evidences were estimated using standard variational Bayesian techniques^69^. Finally, Bayesian model comparison between the N and NH models was performed to compute the corresponding Bayes factors.

### Histology procedures

For standard immunohistochemistry, animals were deeply anesthetized with isoflurane overdose and transcardially perfused with phosphate-buffered saline (PBS) followed by ice-cold 4% paraformaldehyde in PBS. Brains were extracted, post-fixed overnight at 4 °C, and sectioned the following day. Free floating sections (50 μm thick) were cut using a vibratome (Leica VT1200 S, Leica Microsystems GmbH, Wetzlar, Germany). Expression of mCherry was assessed by two-night incubation with a primary anti-mCherry antibody (1:500 dilution, ab205402, Abcam, Cambridge, UK), followed by a three-hours incubation with the secondary antibody Anti-Rabbit Rhodamine Red (1:200 dilution, 711-295-152, Jackson ImmunoResearch, West Grove, PA). Stained brain sections were mounted on glass slides with Invitrogen ProLong Gold Antifade (Fisher Scientific, Waltham, MA) and protected with a coverslip. Fluorescence imaging was performed using a confocal microscope (LSM 900, Zeiss, Thornwood, NY).

For *in-situ* RNA staining with hybridization chain reaction (HCR) amplification^70^, animals were perfused and fixed as for immunohistochemistry, and then processed using the HCR Gold RNA-FISH kit protocol from Molecular Instruments (Los Angeles, CA). Briefly, 50 μm free-floating coronal sections were sliced using a cryostat (Leica Biosystems). Sections were washed with diethyl pyrocarbonate (DEPC)-treated PBS for 3 min each. The sections were then treated with 5% sodium dodecylsulfate in DEPC-PBS for 45 min at room temperature. After rinsing in 2× saline-sodium citrate (SSC) buffer, the sections were incubated in 2× SSC for 15 min. The sections were then incubated in probe hybridization buffer for 30 min at 37 °C for 30 min, followed by incubation with a probe for VGAT (Molecular Instruments) overnight at 37°C. After washing in HCR probe wash buffer (four times for 15 min at 37 °C), the sections were rinsed again in 2× SSC (twice for 5 min) and incubated in HCR Gold Amplifier Buffer (Molecular Instruments) for 30 min at room temperature. The sections were then incubated for 24 hours at room temperature with appropriate RNA hairpins conjugated with Alexa Fluor (denatured and snap-cooled according to the protocol) to visualize hybridization signals. Sections were then washed with HCR Gold Amplifier Wash Buffer (four times for 15 min).

For subsequent visualization of NOSTIC, sections incubated overnight with a primary anti-mCherry antibody (1:500 dilution, ab205402, Abcam, Cambridge, UK), followed by a three-hour incubation with anti-rabbit Alexa Fluor 647 (1:500 dilution, A-31573, Invitrogen, Carlsbad, CA). Stained brain sections were mounted on glass slides with Invitrogen ProLong Gold Antifade (Fisher Scientific, Waltham, MA) and protected with a coverslip. Fluorescence imaging was again performed using a confocal microscope (LSM 900, Zeiss, Thornwood, NY).

### Histological data analysis

Coronal brain slice images acquired using confocal microscopy were exported as TIFF files. For atlas registration, images were downsampled to below 16 megapixels using the Pillow (https://pypi.org/project/pillow/) and OpenCV (https://pypi.org/project/opencv-python/) packages in Python. Downsampled images were registered to the Waxholm Space Rat Brain Atlas version 4^71^ using the DeepSlice rat model^72^, followed by manual refinement in QuickNII^73^. Registration generated three affine transformation vectors that defined the spatial correspondence between each histological section and the standardized atlas coordinate system. Atlas reference slices (< 512 × 512 pixels) were upsampled to match the original image resolution to enable precise delineation of ROIs.

Cell segmentation within each ROI was performed using a CellPose-SAM model^74^ fine-tuned on ten manually annotated images from the dataset. Cell density for each ROI in cells/mm^3^ was defined as *C/NV*, where *C* is the total cell count within the ROI, *N* is the number of atlas voxels encompassing the ROI, and *V* is the volume of each voxel (∼60,000 µm³). For identification of inhibitory neurons, cells positive for each marker (mCherry and VGAT) were segmented independently using finetuned CellPose-SAM models. Co-localization was assessed by calculating the spatial overlap between segmented cell boundaries across the two channels. Putative cells exhibiting greater than 10% overlap, defined as the intersection area divided by the smaller of the two segmented cell areas, were classified as double-positive inhibitory neurons.

## Code availability

Data processing scripts used in analyses presented here are available from the corresponding author upon request.

## Data availability

Source data is provided with this manuscript.

---

**Supplementary Table 1.**
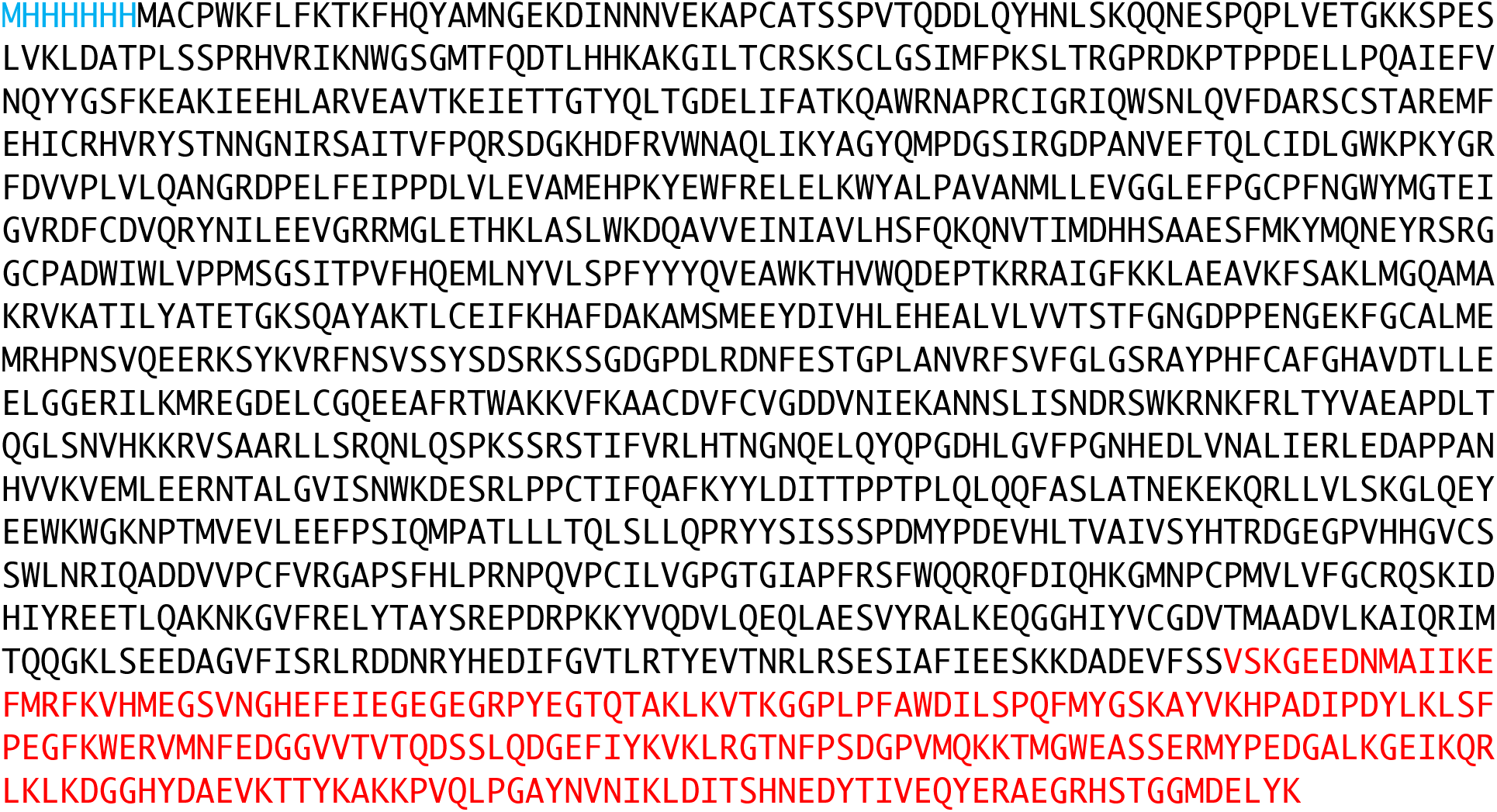
Protein sequence of the NOSTIC-mCherry fusion protein. Sequence of the NOSTIC-mCherry protein encoded by HSV-NOSTIC. The mCherry sequence is color-coded in red and an N-terminal histidine tag is color coded in blue.

**Supplementary Figure 1.**
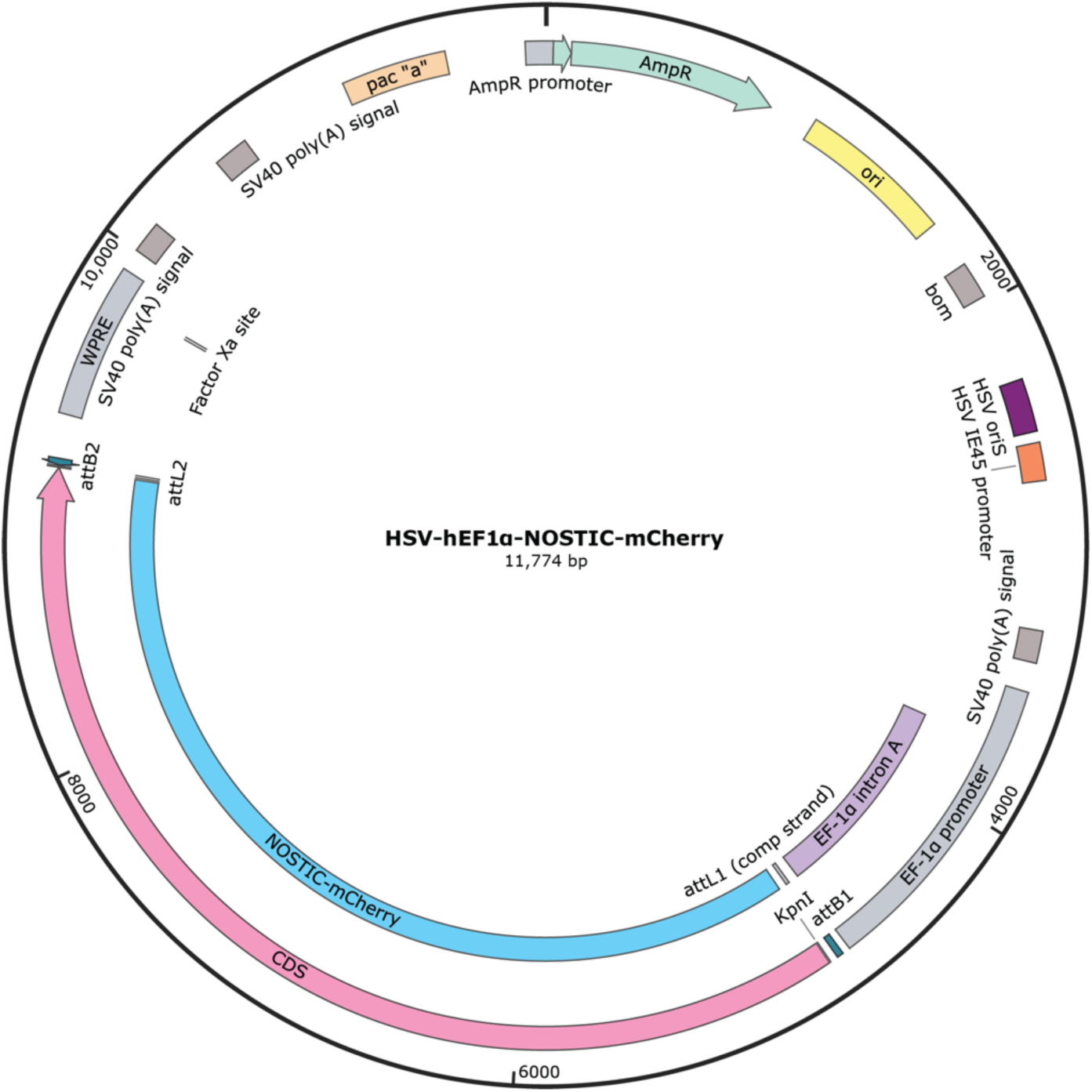
Design of the HSV-NOSTIC construct. Plasmid map indicating major genetic elements in the HSV-NOSTIC construct used throught this study.

**Supplementary Figure 2.**
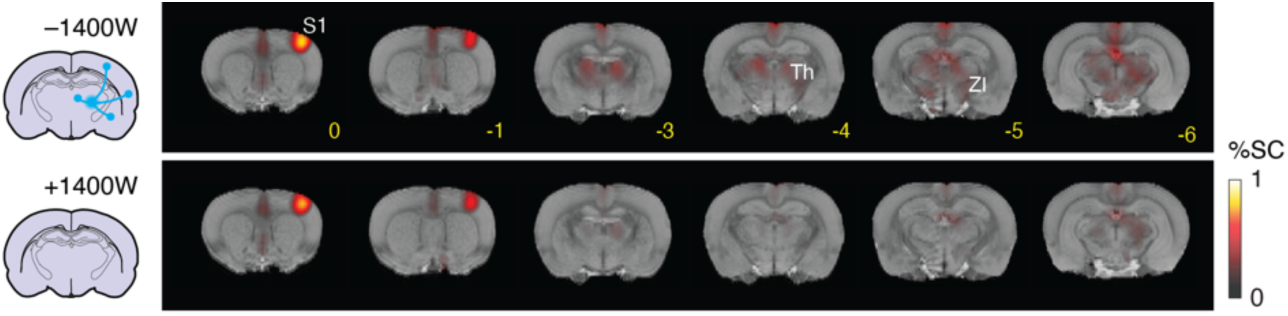
Activation maps from animals treated with HSV-mCherry. Activation maps showing response to forelimb stimulation of animals treated with HSV-mCherry in POm. Color overlays indicate mean percent signal changes from 6 animals in the absence (top) and presence (bottom) of 1400W. Signal change shown for voxels with significant stimulus responses (FDR-corrected *F*-test *p* ≤ 0.05, *n* = 8). Bregma coordinates in yellow.

**Supplementary Figure 3.**
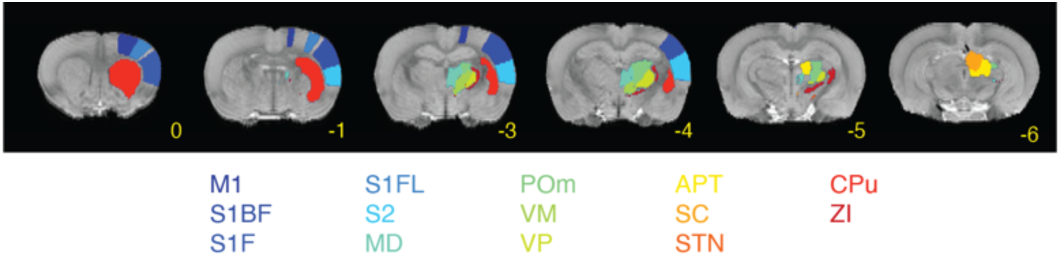
Definition of ROIs used for regional analyses. Brain regions used for ROI analyses throughout this study are defined by colored areas corresponding to the color-coded labels shown at bottom. Unilateral regions are shown here; analogous regions in the opposite hemisphere are symmetrically related.

**Supplementary Figure 4.**
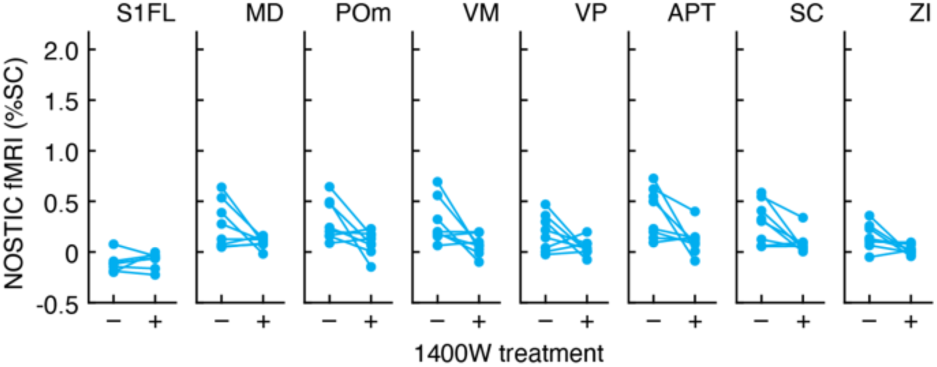
1400W dependence of fMRI responses contralateral to HSV-NOSTIC injection sites. Amplitudes of fMRI signal changes are shown in the absence and presence of 1400W for regions contralateral to HSV-NOSTIC injection sites and ipsilateral to forelimb stimulation as in Figure 1. 1400W-dependent signal differences in PoM, VM, APT, SC and ZI are significant with paired *t*-test *p* ≤ 0.048, *n* = 8.

**Supplementary Figure 5.**
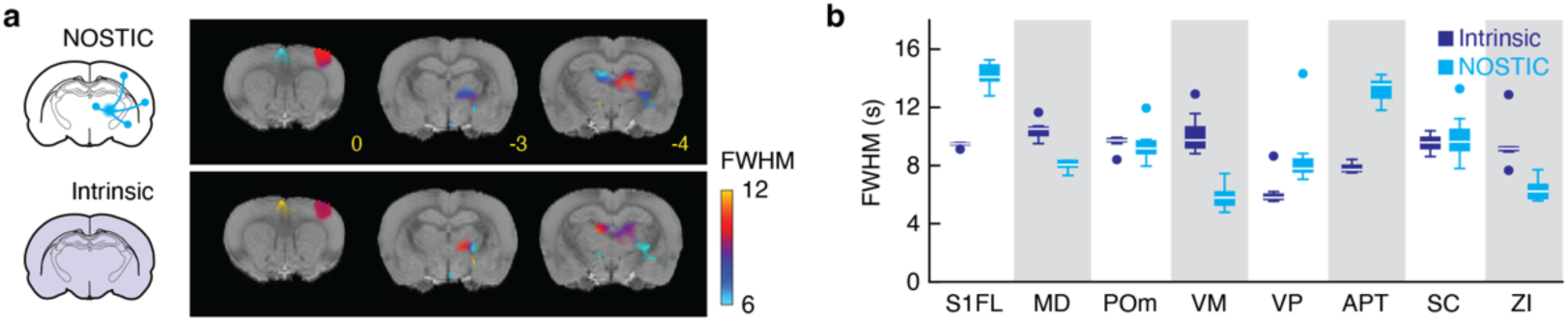
Duration of NOSTIC and intrinsic responses to stimulation. **a**, Maps showing duration (FWHM) of NOSTIC and intrinsic fMRI responses in selected coronal brain sections, as computed from gamma function curve fits to voxel-level time courses. Data presented for voxels with significant NOSTIC signal (FDR-corrected *F*-test *p* ≤ 0.05). **b**, FWHM values observed in regions of interest. Box plot denotes median (center line), first and third quartiles (boxes), full range (whiskers), and outliers (dots).

**Supplementary Figure 6.**
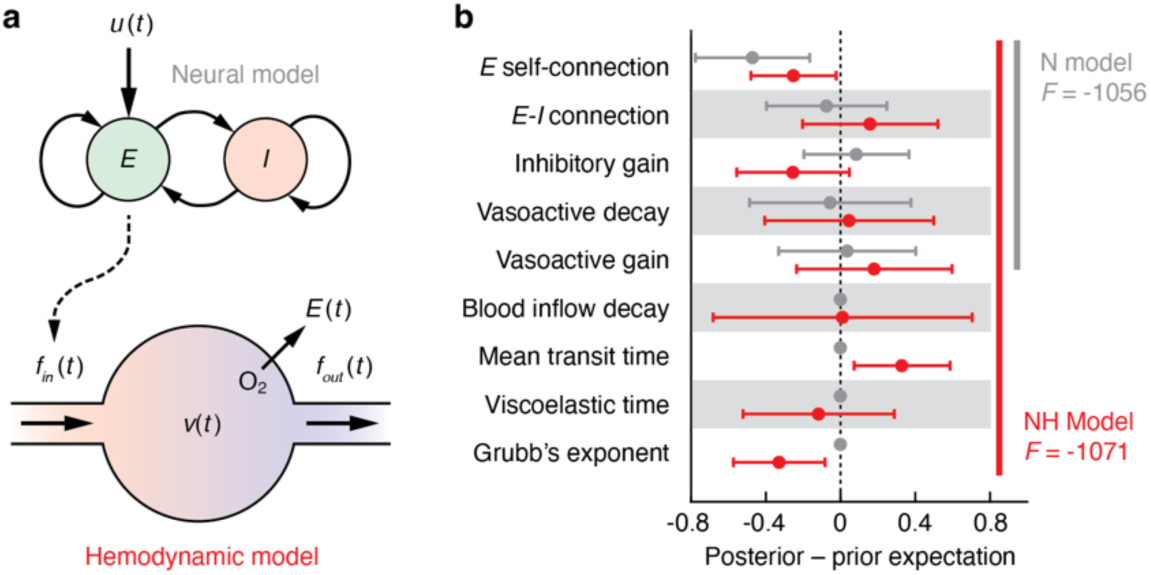
Hemodynamic and neural factors for NOSTIC response modeling. **a**, Diagram showing key elements of N and NH models. The top portion of the diagram illustrates driving external input u(t), and local network activity represented by self-connected and reciprocally coupled excitatory (*E*) and inhibitory (*I*) activity components. These elements in turn provide input (dashed arrow) to the hemodynamic model depicted at bottom. Excitatory activity specifically drives blood inflow into local vasculature, *f_in_*(*t*), which in turn influences the local blood volume *v*(*t*), outflow *f_out_*(*t*), and oxygen extraction fraction *E*(*t*) via a set of viscoelastic parameters, following the formalisms of ref. 34. In the N model but not the NH model, the hemodynamic model parameters are fixed. b, Posterior minus prior expectation values obtained for specific parameters listed at left, under the N (gray) and NH (red) models, after fitting to the 1400W-dependent fMRI data of Figure 1. Expectation ranges overlap closely for the two models. The variational free-energies *F*, upper-bounding log-model evidences, are noted at right for both models. Higher values indicate more accurate or less complex models.

**Supplementary Figure 7.**
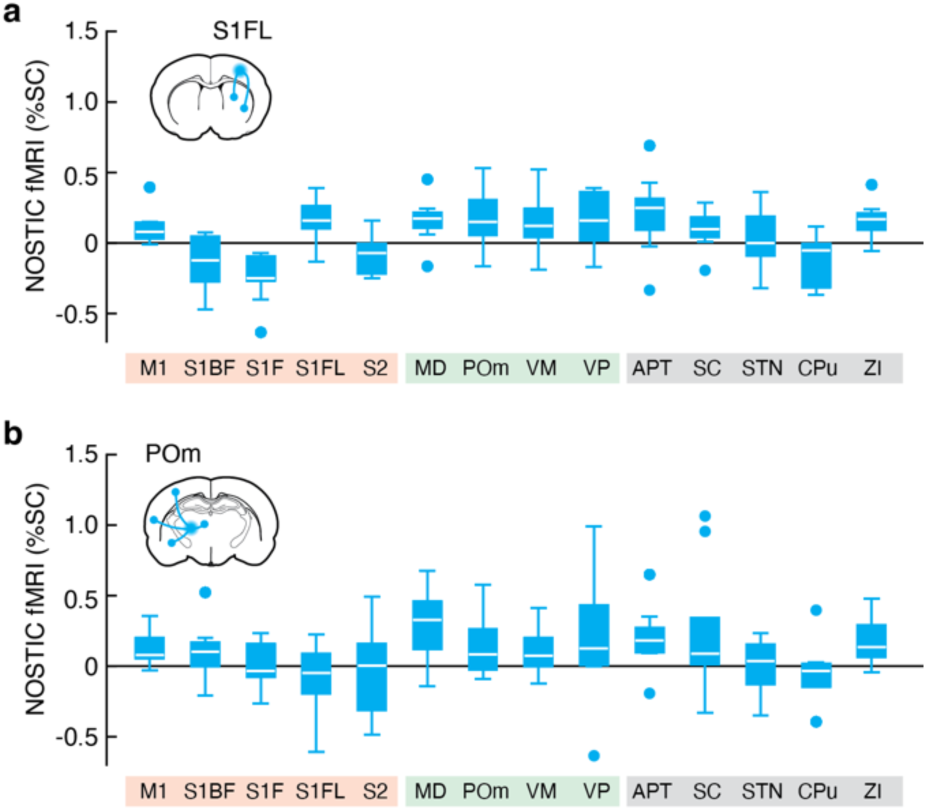
NOSTIC responses to forelimb stimulation. **a**, NOSTIC response amplitudes analogous to those in Figure 1f were determined form experiments in animals treated with HSV-NOSTIC injections into S1FL, contralateral to the stimulated forelimb. Responses in M1, S1BF, S1F,S1FL,MD,POm,VP and ZI are significant with paired *t*-test *p* ≤ 0.48 (*n* = 9). **b**, NOSTIC amplitudes were observed in animals injected with HSV-NOSTIC in POm ipsilateral to the stimulated limb. Responses in MD and APT are significant with paired *t*-test *p* ≤ 0.03 (*n* = 8). For both panels, labels on the horizontal axis denote cortical, thalamic, and other subcortical ROIs. Box plots denote median (center line), first and third quartiles (boxes), full range (whiskers), and outliers (dots).

**Supplementary Figure 8.**
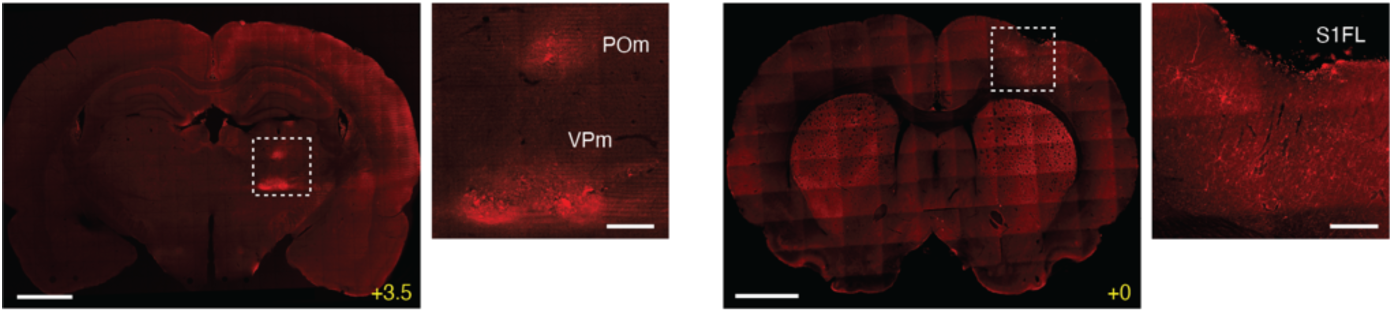
Visualization of NOSTIC expression animals treated virus in S1FL. Representative histological data showing expression in cortical and subcortical regions of animals injected with HSV-NOSTIC in S1FL. Scale bars: 2 mm (full slices), 400 µm (closeups corresponding to dashed boxes).

**Supplementary Figure 9.**
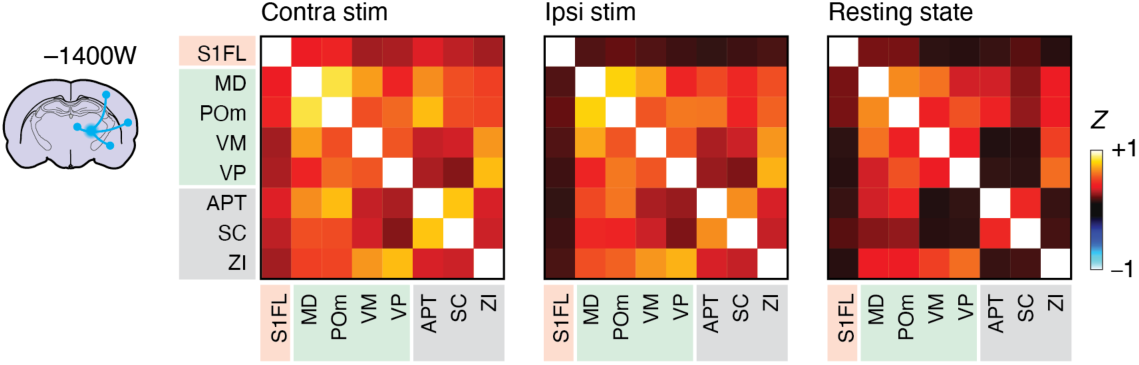
FC matrices from NOSTIC-treated animals in the absence of 1400W. Z values for FC matrices in the –1400W condition, reflecting superposition of intrinsic and NOSTIC signals, during contralateral forelimb stimulation (left), ipsilateral stimulation (middle), and resting state (right) conditions. Green outlines denote significance of individual matrix elements with t-test *p* ≤ 0.05 (*n* = 8 for contra and ipsi stim, *n* = 9 for resting).

**Supplementary Figure 10.**
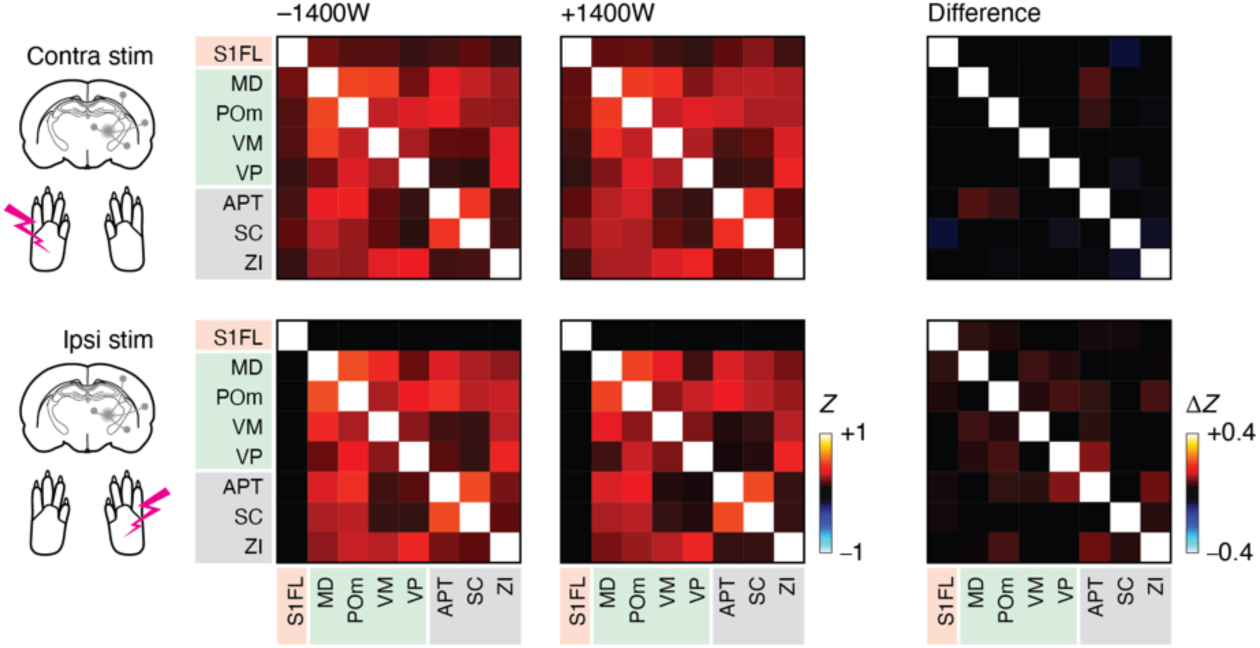
FC data from animals treated with HSV-mCherry. FC matrices computed from animals treated with HSV-NOSTIC in POm and stimulated contralateral (top) or ipsilateral (bottom) to the viral injection sites. Z matrices obtained in the –1400W (left) and +1400W (middle) conditions are virtually indistinguishable, leading to negligible Δ*Z* values under both stimulation conditions in the matrices at right.

**Supplementary Figure 11.**
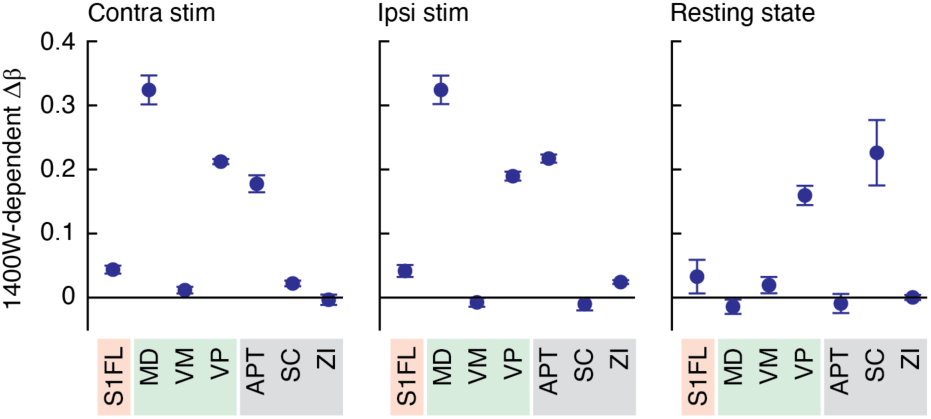
Regression models for intrinsic POm signal under three conditions. Post-1400W intrinsic signal in POm was modeled as a linear combination of fMRI signals from regions that provide ipsilateral NOSTIC-detectable input to POm. Graphs show regression coefficients calculated from data obtained during contralateral fore-limb stimulation (left), ipsilateral stimulation (middle), and rest (right). Dots: means, error bars: SEM over 8 or 9 jackknifed animals.

